# Single-Cell RNA-seq of Heart Reveals Intercellular Communication Drivers of Myocardial Fibrosis in Diabetic Mice

**DOI:** 10.1101/2022.06.29.498129

**Authors:** Wei Li, Xinqi Lou, Yingjie Zha, Jun Zha, Lei Hong, Zhanli Xie, Shudi Yang, Chen Wang, Jianzhong An, Zhenhao Zhang, Shigang Qiao

## Abstract

Diabetes-induced cardiomyopathy is characterized by myocardial fibrosis as a main pathology. In-depth study of cardiac heterogeneity and cell-to-cell interactions will help to reveal the pathogenesis of diabetic myocardial fibrosis and provide potential targets for the treatment of this disease. Here, we insighted into the intercellular communication drivers underlying myocardial fibrosis in mouse heart with high-fat-diet (HFD)/streptozotocin (STZ)-induced diabetes at single-cell resolution. Intercellular and protein-protein interaction networks of fibroblasts and macrophages, endothelial cells, as well as fibroblasts and epicardial cells reveal critical changes in ligand-receptor interactions such as Pdgf(s)-Pdgfra and Efemp1-Egfr, which promote the development of a profibrotic microenvironment during diabetes progression and confirm that specific inhibition of Pdgfra axis can significantly improve diabetic myocardial fibrosis. We further identified the phenotypically distinct Hrc^hi^ and Postn^hi^ fibroblast subpopulations that are associated with pathological extracellular matrix remodeling, of which Hrc^hi^ fibroblasts are the most profibrogenic under diabetic conditions. Finally, we validated the role of Itgb1 hub gene mediated intercellular communication drivers of diabetic myocardial fibrosis in Hrc^hi^ fibroblasts, and confirmed the result by AAV9-mediated Itgb1 knockdown in the heart of diabetic mice. In summary, cardiac cell mapping provides novel insights into intercellular communication drivers underlying pathological extracellular matrix remodeling during diabetic myocardial fibrosis.

## Introduction

Diabetes is the third cause of threatening human health, approximately 537 million adults are living with diabetes, of which type 2 diabetes patients account for more than 90%. Cardiac complications are the most common causes of death and disability associated with diabetes. As the key initiating factor of diabetic cardiomyopathy, hyperglycemia can prevent optimal utilization of glucose by the cardiomyocytes and leads to myocardial fibrosis. Myocardial fibrosis is characterized by the increase in extracellular matrix proteins, deposition of interstitial collagen, disarrangement of cardiomyocytes and the remodeling of cardiac structure *(Russo and Frangogiannis, 2016; Jia et al., 2018)*. Since adult mammalian cardiomyocytes are virtually incapable of regeneration, the most extensive extracellular matrix remodeling and fibrosis of the heart occurs in diseases caused by acute cardiomyocyte death *(Kong et al., 2014)*. Understanding the mechanisms responsible for myocardial fibrosis is critical to develop anti-fibrotic therapy strategies for diabetic patients.

Mammalian hearts consist of many cell types, including cardiomyocytes, macrophages, fibroblasts and endothelial cells, etc. *(Banerjee et al., 2007; Litviňuková et al., 2020)*. Cell-to-cell communication is a fundamental feature of adult complex organs. These different types of cells communicate through interactions of ligand-receptor, where a ligand may be secreted and bind to a receptor, or through the fusion of two adjacent interacting cell membranes *(Ramilowski et al., 2015)*. The maintenance of heart homeostasis depends on the intercellular communication *(Ramilowski et al., 2015)*. Many ligand-receptor signaling patterns have been found between the cardiac cells, suggesting the critical role of intercellular communication in many pathophysiological processes. Therefore, intercellular communication has become a powerful therapeutic target for preventing or reversing some of the damaging consequences of diabetic myocardial fibrosis by maintaining fine-tuned intercellular communication among different cardiac cells. Despite these, the effect of diabetes on cardiac intercellular communication and myocardial fibrosis remains poorly understood.

Single-cell RNA sequencing (scRNA-seq) is a feasible strategy to study the cellular heterogeneity of any organ since it allows transcriptomic profiling of individual cells *(Butler et al., 2018; Gladka et al., 2018; Litviňuková et al., 2020; McLellan et al., 2020)*. Recent scRNA-seq of many tissues has revealed cellular heterogeneity and novel intercellular crosstalk among different cell types. In this study, we developed a diabetic mouse model through a HFD combined with STZ administration, and analyzed cell populations in mouse heart by scRNA-seq on a 10x genomics platform.

We profiled 32,585 single cardiac cell transcriptomes across 6 healthy controls and 6 diabetic mice, and identified the transcriptional alterations of all cardiac cells, enrichment of signaling pathways involved in myocardial fibrosis, altered ligand-receptor interactions described as Pdgf(s)-Pdgfra and Efemp1-Egfr between fibroblasts and other cardiac cells, and cellular subpopulations associated with diabetic myocardial fibrosis. Furthermore, a specific Hrc^hi^ fibroblast subcluster as well as intercellular communication drivers of Itgb1 hub gene mediated myocardial fibrosis were identified. These results suggest that cardiac intercellular communication plays a critical role in diabetic myocardial fibrosis and specific targeting of Hrc^hi^ fibroblasts may be a potential therapeutic target for diabetic myocardial fibrosis.

## Results

### Single-cell profile of heart in diabetic mice

Conventional single-cell RNA-seq cannot encompass all the cells in the rodent myocardium for subsequent deep sequencing *(Skelly et al., 2018; Forte et al., 2020)*. Therefore, we isolated the nuclear fractions of all cardiac cells to assess the heterogeneity of cell populations and the changes of transcriptional profile in response to pathology of HFD/STZ-induced diabetes (Figure 1-Figure supplement 1A) *(Grindberg et al., 2013; Lake et al., 2019)*. We classified 32585 cardiac cells from 6 healthy controls (16490 cells) and 6 HFD/STZ-induced diabetic mice (16095 cells) into 25 transcriptionally distinct pre-clusters exhibiting highly consistent expression patterns across individual mice (Figure 1-Figure supplement 1B and Supplementary file 1), and identified 14 populations (Figure 1A) based on cell-specific markers and the significantly enriched genes. The cell populations included fibroblasts (Pdgfra, Pcdh9, Bmper), endothelial cells (Pecam1, Ccdc85a, Btnl9), macrophages (Fcgr1, F13a1, Adgre1), pericytes (Pdgfrb, Vtn, Trpc3), adipocytes (Adipoq, Plin1, Tshr), cardiomyocytes (Ttn, Mhrt, Myh6), smooth muscle cells (Acta2, Myh11, Cdh6), endocardial cells (Npr3, Tmem108, Plvap), epicardial cells (Msln, Pcdh15, Muc16), schwan cells (Plp1, Gfra3), and other immune cell populations (T cells, Monocytes, B cells) (Figure 1B). Based on the cell types, markers, and relative proportions, we can conclude that our data are robust and consistent with previous 10× single nucleus data from mice heart. *(McLellan et al., 2020)*. Further examination of the established marker genes in each cardiac cell population revealed the presence of a wide range of cell types in all hearts (Figure 1C and D).

**Figure 1.**
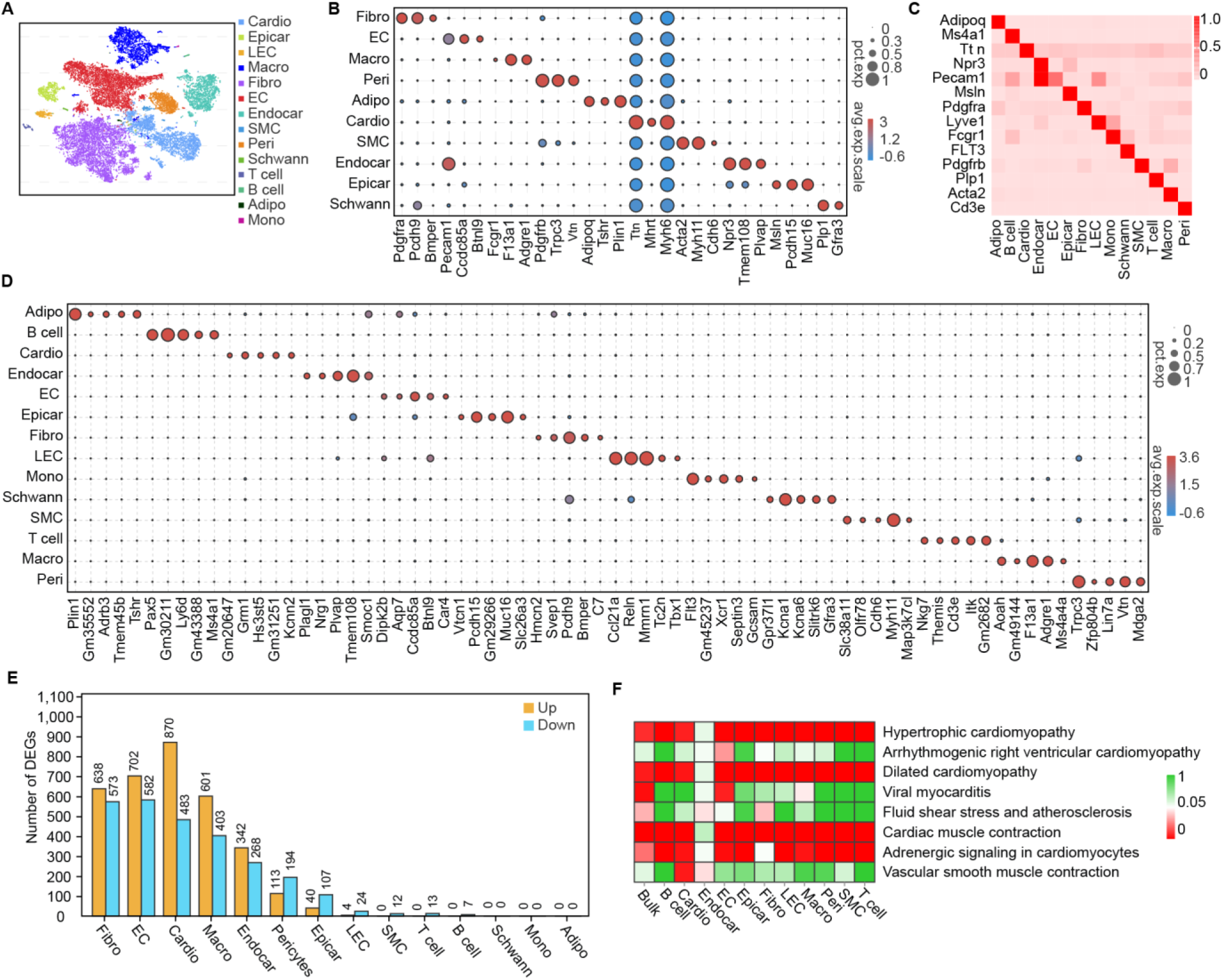
Single-cell profile of heart in diabetic mice. (A) t-SNE projection of all mouse cardiac cells (n = 16490 cardiac cells from 6 control mice and n = 16095 cardiac cells from 6 diabetic mice). (B) The marker genes defining each type of cell cluster in A are listed. The circle size illustrates the proportion of cells expressing each transcript within each group. The dot color represents the expression level within each population. Color scale: red, high-expressive level; blue, high-expressive level. (C) Heat map shows the canonical cell markers of major cardiac cell populations. (D) Dot plot represents the top 5 distinct genes for each cell population. (E) Lollipop plot shows number of high and low expressed genes in HFD/STZ-treated mouse heart cells relative to controls (2-sided Wilcoxon rank-sum test, FDR≤0.05, log2FC≥ 0.36). (F) Heatmap shows enriched KEGG pathways associated with cardiovascular diseases and circulatory system in each cardiac cell population in the diabetic group. Color scale: red, low FDR; green, high FDR (2-sided Wilcoxon rank-sum test, FDR≤ 0.05). Adipo, adipocytes; Cardio, cardiomyocytes; Endocar, endocardial cells; EC, endothelial cells; Epicar, epicardial cells; Fibro, fibroblasts; LEC, lymphatic ECs; Mono, monocytes; Schwann, schwann cells; SMC, smooth muscle cells; Macro, macrophages; Peri, pericytes; FDR, false discovery rate. The details of 25 transcriptionally distinct pre-clusters with highly consistent expression patterns across individual mouse heart are listed in *Supplementary file 1*. Detailed genes of significant transcriptomic changes in cardiac populations are listed in *Supplementary file 2*. The details of top 10 upregulated genes in cardiac populations are listed in *Supplementary file 3*. This paper includes the following figure supplement(s) for Figure 1. **Figure supplement 1.** Experimental design and cell type characterization. **Figure supplement 2.** The top 10 upregulated genes during pathology of HFD/STZ-induced diabetes within each cell population. **Figure supplement 3.** Top 30 enriched KEGG pathways in HFD/STZ-treated mouse fibroblasts. **Figure supplement 4.** Top 30 enriched KEGG pathways in HFD/STZ-treated mouse cardiomyocytes, endothelial cells, endocardial cells, and macrophages.

**Figure 1-Figure supplement 1.**
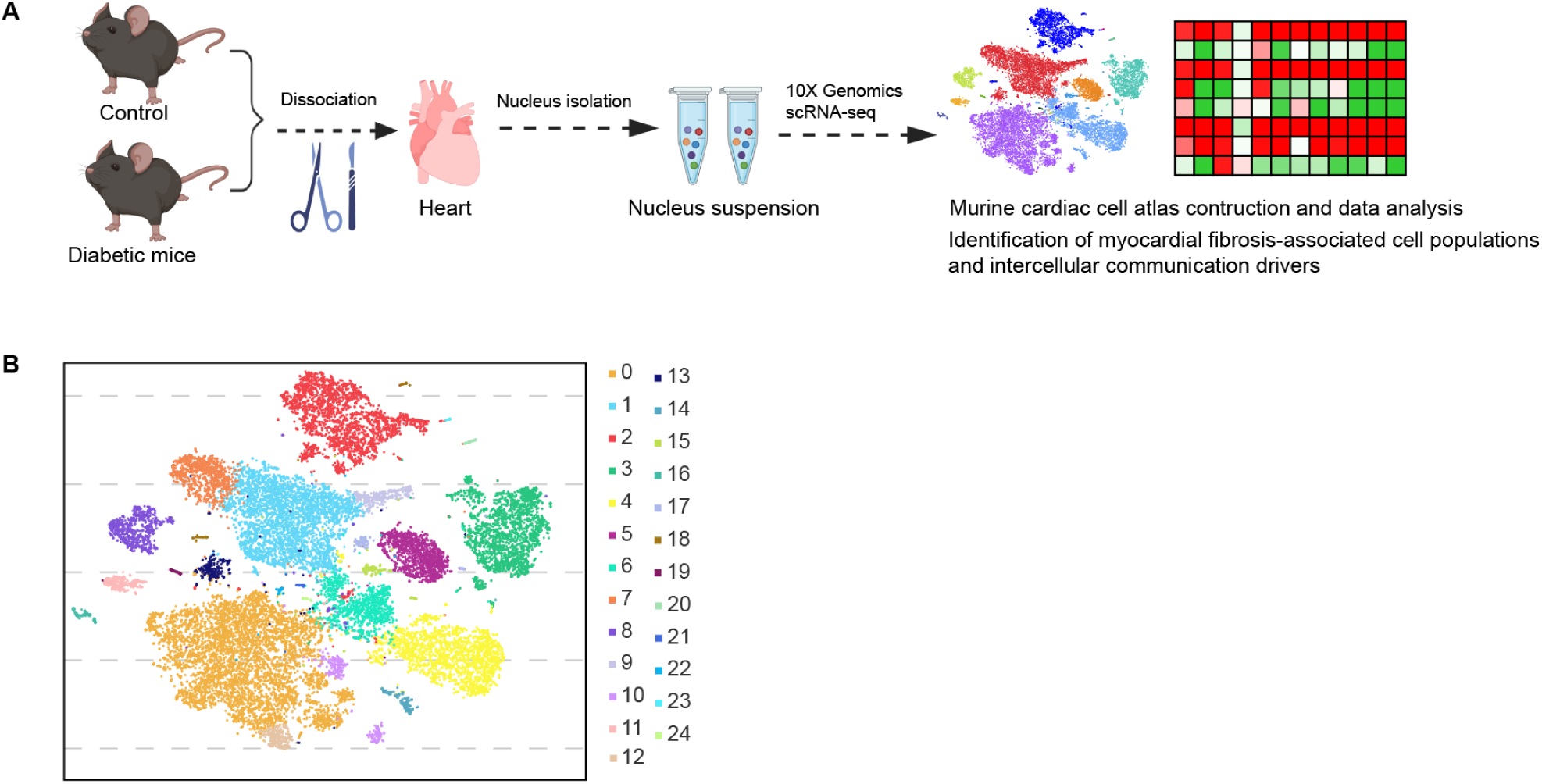
Experimental design and cell type characterization. (A) Schematic overview of experimental design. (B) t-SNE projection of all pre-clustered cells (n = 16490 cardiac cells from 6 control mice and n = 16095 cardiac cells from 6 diabetic mice).

**Figure 1-Figure supplement 2.**
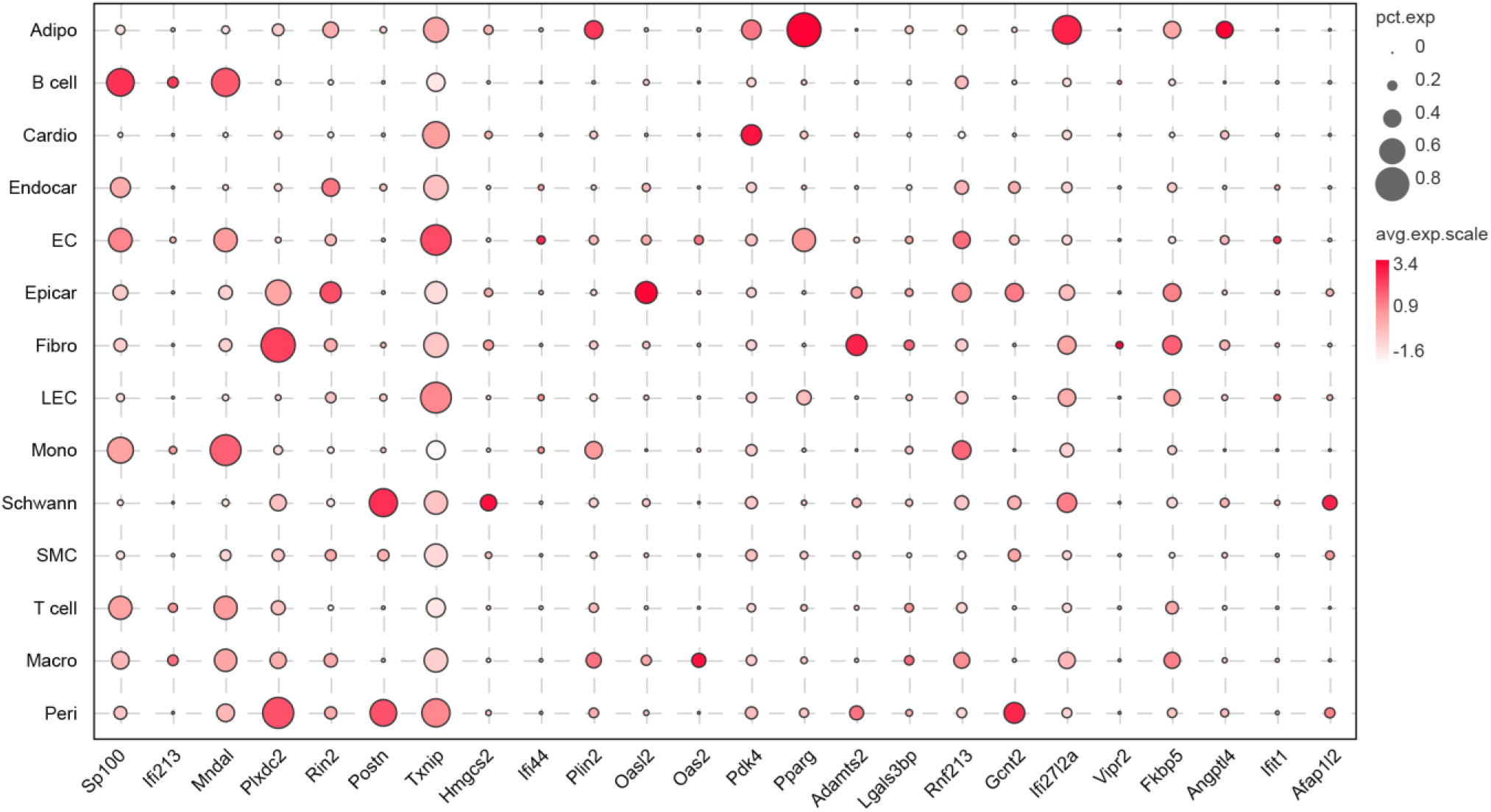
Dot plot represents the top 10 upregulated genes during pathology of HFD/STZ-induced diabetes within each cell population. The circle size illustrates the proportion of cells within each transcript, the dot color indicates the relative average expression level of the gene.

**Figure1-Figure supplement 3.**
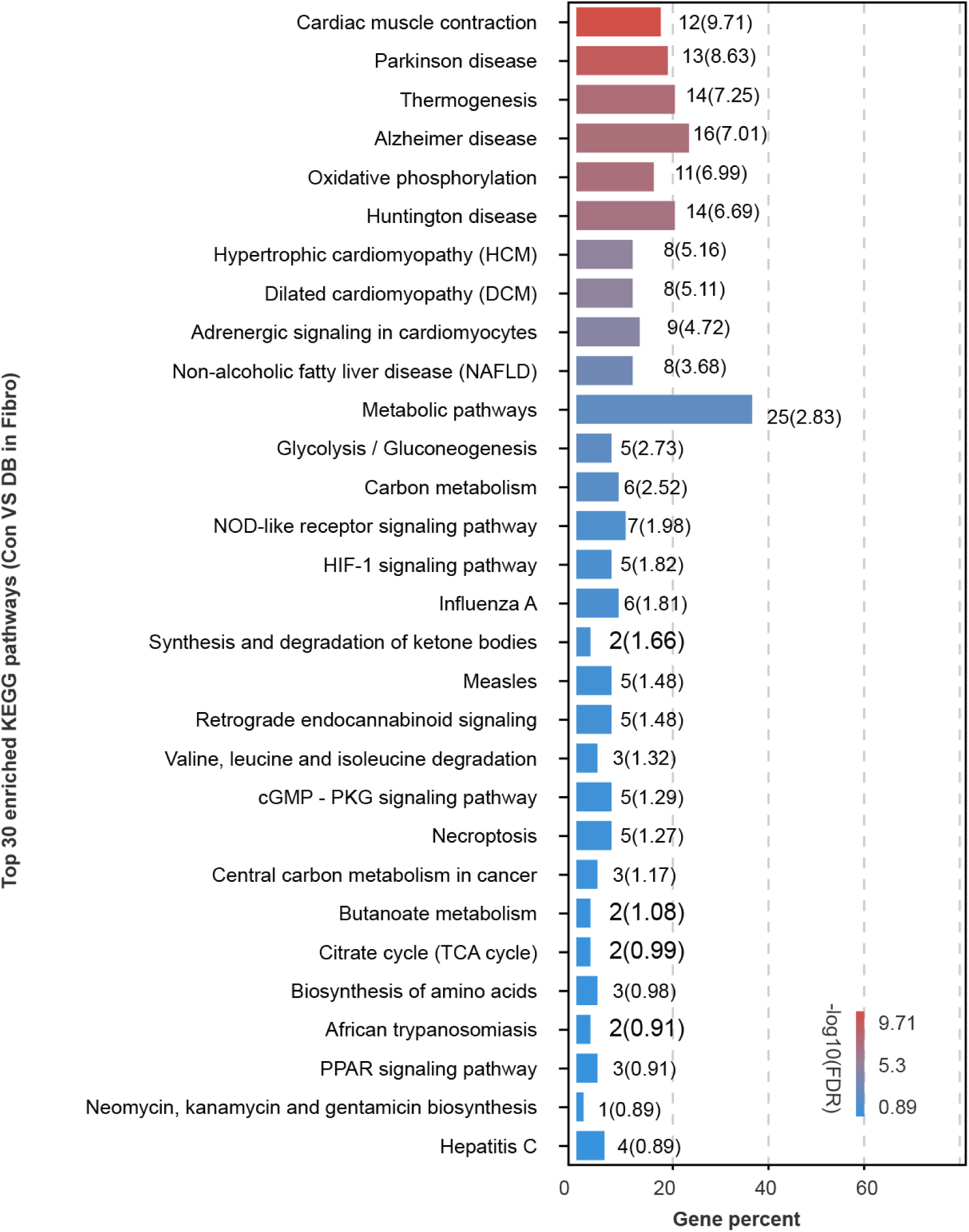
Plot shows top 30 enriched KEGG pathways in HFD/STZ-treated mouse fibroblasts relative to control (2-sided Wilcoxon rank-sum test, FDR≤0.05). Color scale: red, high-log10(FDR) value; green, low-log10(FDR) value.

**Figure 1-Figure supplement 4.**
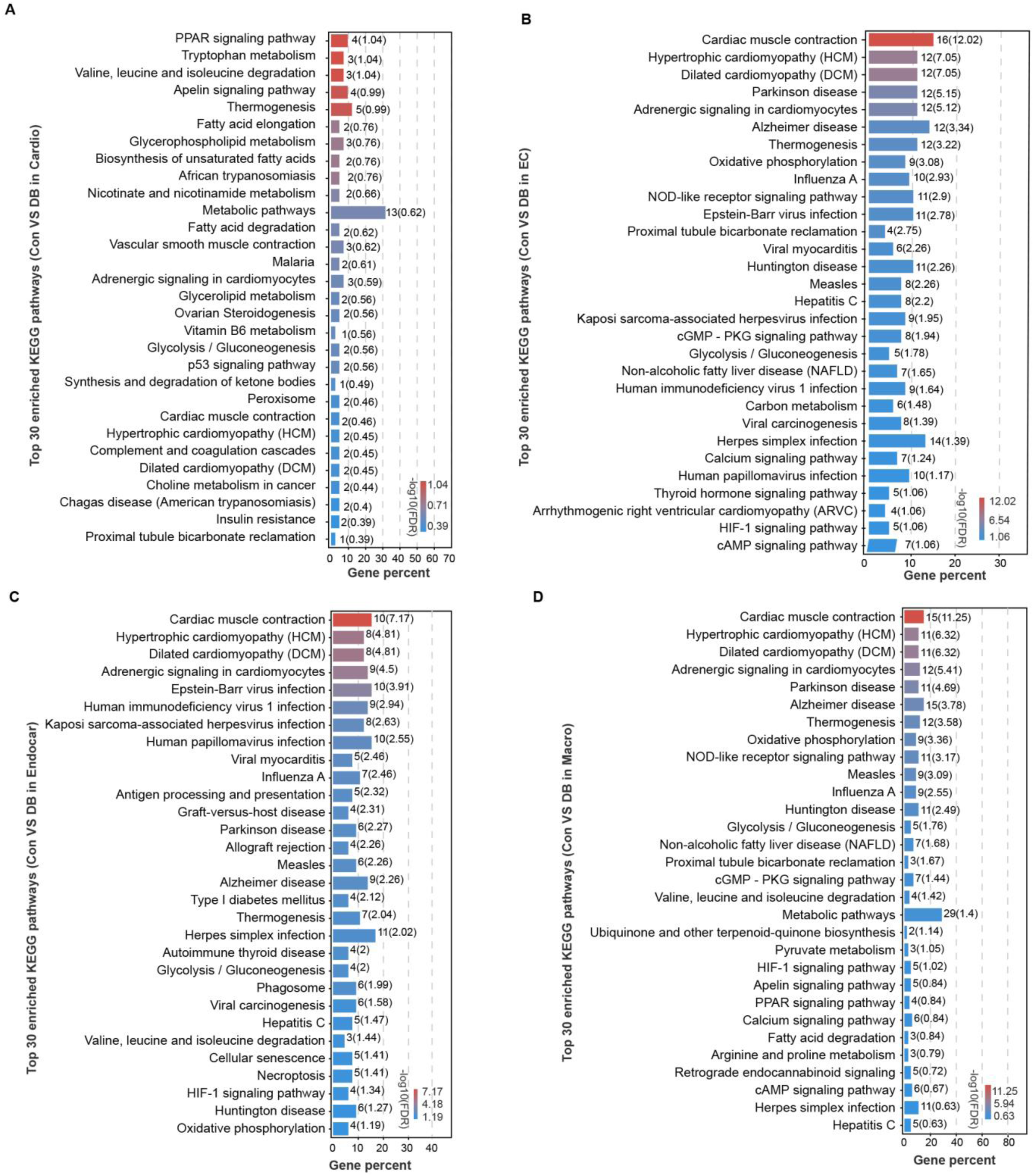
Plot shows top 30 enriched KEGG pathways in HFD/STZ-treated mouse cardiomyocytes (A), endothelial cells (B), endocardial cells (C), and macrophages (D) relative to control (2-sided Wilcoxon rank-sum test, FDR≤ 0.05). Color scale: red, high-log10(FDR) value; green, low-log10(FDR) value.

HFD/STZ-treatment induced significant transcriptomic changes in most cardiac populations, especially in fibroblasts, endothelial cells, endocardial cells, cardiomyocytes and macrophages (Figure 1E, Supplementary file 2, 2-sided Wilcoxon rank-sum test, FDR≤ 0.05, log2FC≥0.36). Analyzing the top 10 upregulated genes during pathology of HFD/STZ-induced diabetes within each cell population, we found that some of the top 10 upregulated genes show a noticeable increase in expression across many different cell types, even though some are primarily expressed in only one cell type (Figure 1-Figure supplement 2 and Supplementary file 3). Among the top genes upregulated in response to HFD/STZ-induced diabetes within each cell population were transcripts for PDK4, Angptl4, Txnip, Postn, Hmgcs2, and Ucp3, several of which have been previously involved in heart failure. *(Yoshioka et al., 2007; Lang et al., 2018; Sheeran et al., 2019)* KEGG (Kyoto Encyclopedia of Genes and Genomes) analysis of the upregulated genes revealed significant enrichment of pathways related to cardiovascular diseases and circulatory system in most cardiac cell types (Figure 1F, 2-sided Wilcoxon rank-sum test, FDR≤ 0.05), and the top 30 KEGG pathways within some cardiac cell populations were relevant to dilated cardiomyopathy, cardiac muscle contraction, and hypertrophic cardiomyopathy, (Figure 1-Figure supplement 3, Figure 1-Figure supplement 4A-D, 2-sided Wilcoxon rank-sum test, FDR≤0.05). These results indicated a significantly increased risk of cardiovascular diseases in the diabetic setting.

**Figure 2.**
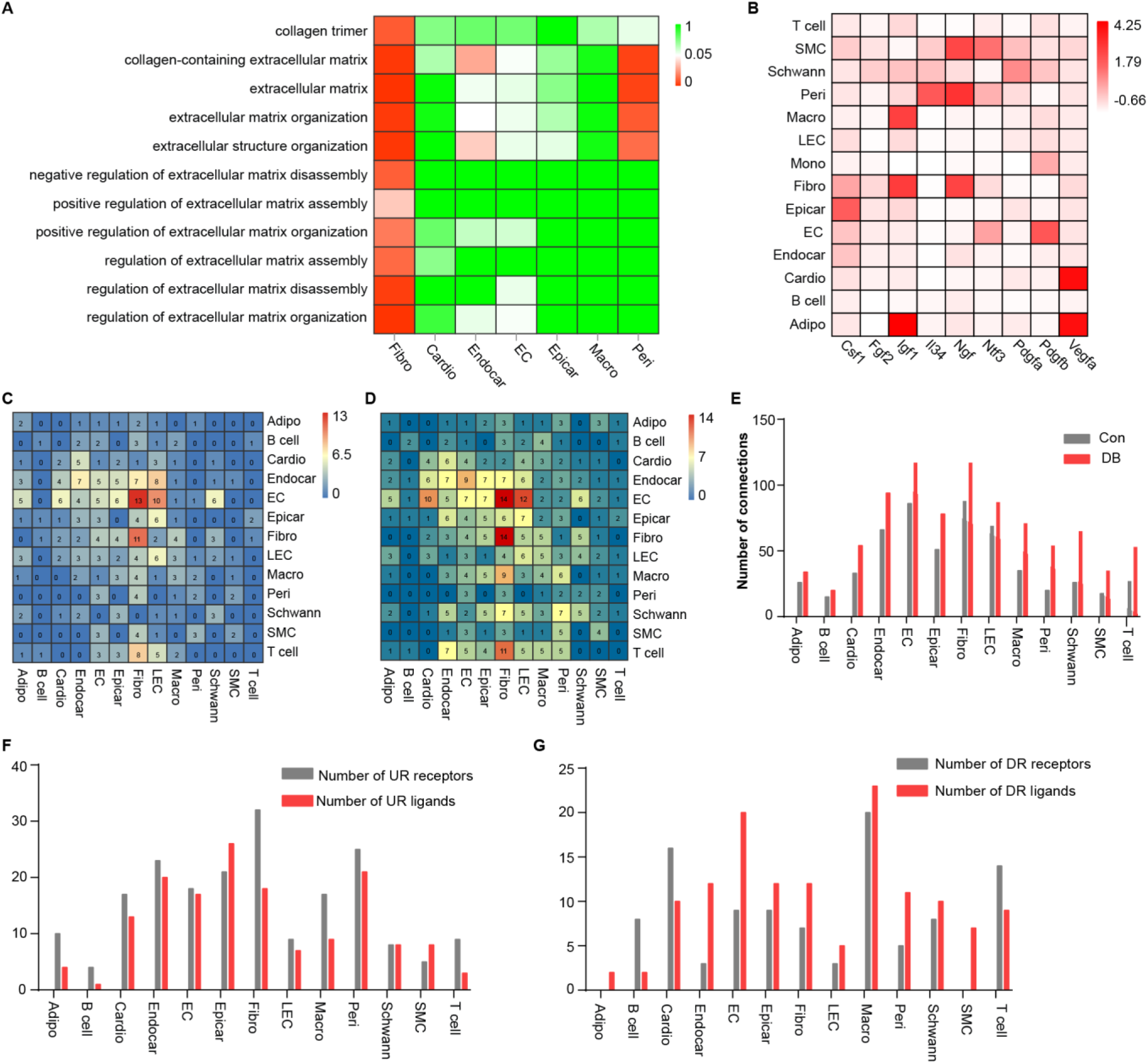
Comparison analysis of the communications between cardiac cells during HFD/STZ-induced diabetes. (A) Heatmap shows enriched GO terms associated with extracellular matrix remodeling and myocardial fibrosis in major cardiac cell populations in the diabetic group (2-sided Wilcoxon rank-sum test, FDR≤0.05). Color scale: red, low FDR; green, high FDR. (B) The relative expression of selected essential growth factors in major cardiac cell types. (C) Heatmap shows the number of ligand–receptor pairs between cardiac cell populations in healthy mice (FDR≤0.01, log2FC≥1). (D) Heatmap shows the number of ligand–receptor pairs between cardiac cell populations in HFD/STZ-induced diabetic mice (FDR≤0.01, log2FC≥1). (E) Bar plot shows total number of connections made by each cell type without (gray bars) and with HFD/STZ treatment (red bars) (2-sided Wilcoxon rank-sum test, FDR ≤0.05, log2FC≥0.36). (F) Bar plot illustrates the number of upregulated receptors and ligands for each population of cardiac cells (2-sided Wilcoxon rank-sum test, FDR≤0.05, log2FC≥0.36). (G) Bar plot shows number of downregulated receptors and ligands for each cardiac cell population. (2-sided Wilcoxon rank-sum test, FDR≤ 0.05, log2FC≥0.36). DB, diabetes. The details of unique differentially-expressed genes (uni-DEGs) in cardiac populations are listed in *Supplementary file 4*. The details of significantly differentially-expressed genes in specific cell populations relative to others in mouse heart are listed in *Supplementary file 5*. Details of cell type-specific receptors in cardiac populations and cell type-specific ligands in cardiac populations are listed in *Supplementary file 6* and *Supplementary file 7*, respectively. The details of relative expression of a selection of essential growth factors across major cardiac cell types are listed in *Supplementary file 8.* The details of the number of ligand-receptor pairs between cardiac cell populations in healthy mice or diabetic mice are listed in *Supplementary file 9* and *Supplementary file 10,* respectively. The details of significant differentially-expressed ligands and receptors for each cell population are listed in *Supplementary file 11* and *Supplementary file 12*, respectively. This paper includes the following figure supplement(s) for Figure 2. **Figure supplement 1.** The expression of receptors and ligands across major cardiac cell types.

**Figure 2-Figure supplement 1.**
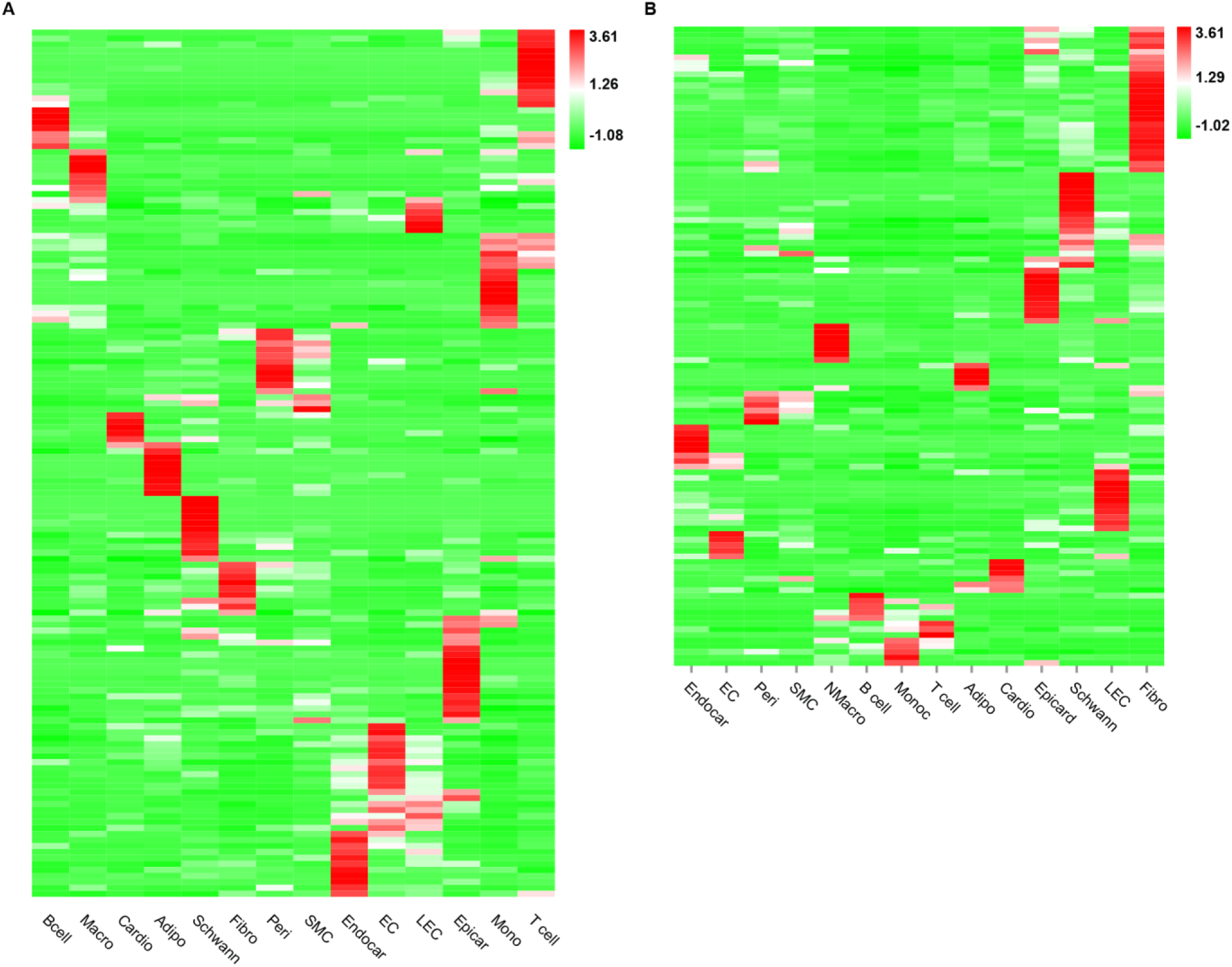
The heatmap shows the relative expression of receptors (A) and ligands (B) across major cardiac cell populations. Red color indicates high expression; green color indicates low expression (FDR≤0.01, log2FC ≥1).

Taken together, scRNA-seq identified distinct cell populations of mouse heart that can help characterize HFD/STZ-induced diabetes-related changes in gene expression and quantify gene-trait associations.

### Effects of HDF/STZ-induced diabetes on cardiac intercellular communication

Cell subpopulations with non-overlapping functions present distinct transcriptomic perturbations in response to pathological stimuli *(Mathys et al., 2019)*. To determine whether distinct cardiac cell populations response heterogeneously to the diabetic stimuli, we compared differentially-expressed genes of all cardiac cell types and identified 2118 unique differentially-expressed genes (uni-DEGs) associated with major cardiac cell types (Supplementary file 4). Most of the uni-DEGs (96.6%) were detected in cardiomyocytes (32.8%), fibroblasts (18.5%), macrophages (17.7%), endothelial cells (19.7%) and endocardial cells (7.9%). The genes most highly expressed in only each cell type were GM20658 (fibroblasts), Ucp3 (cardiomyocytes), Spock2 (endocardial cells), Irf7 (endothelial cells) and Ifi206 (macrophages), of which Ucp3 and Irf7 are involved in heart failure and pathological cardiac hypertrophy *(Jiang et al., 2014; Senatus et al., 2020)*. However, the association of GM20658, Spock2 and Ifi206 with myocardial fibrosis or heart failure has not been reported so far. Gene Ontology (GO) analysis of the uni-DEGs (upregulated) showed that terms associated with collagen fibril organization and extracellular matrix remodeling were enriched in cardiac fibroblasts (Figure 2A, 2-sided Wilcoxon rank-sum test, FDR≤0.05), indicating that fibroblasts are key cellular contributors to extracellular matrix remodeling and cardiac fibrosis.

The proper functioning of metazoans is tightly controlled by the intercellular communication between multiple cell populations, which is based on interactions between secretory ligands and receptors. *(Ramilowski et al., 2015)*. To determine the effect of HFD/STZ-induced diabetes on cardiac intracellular communication, we mapped intercellular connection network of the cardiac cellulome in healthy controls and diabetic mice. Initially, we identified genes that were differentially expressed in specific cell populations in the mouse heart, focusing on those over-expressed in a single cell type, i.e., specific highly-expressed genes, at FDR≤0.01 with a minimum twofold difference in expression (Supplementary file 5). Gene expression patterns for receptors and ligands were found to be cell type specific in the heart secretome genes analyzed by clustering (Figure 2-Figure supplement 1A and B, Supplementary file 6, Supplementary file 7). Analysis of the factors that support the growth of specific cell populations has revealed critical intercellular communication. These include signaling pathways that support the survival of specific cell populations of mouse hearts (Figure 2B, Supplementary file 8). For instance, pericytes and fibroblasts express Il34 and CSF1, respectively (Figure 2B), which communicate through CSF1R and are key factors for macrophage survival and growth. Fibroblasts also express IGF1 and NGF (Figure 2B), which support the growth of endothelial cells, mural cells and neurons *(Glebova and Ginty, 2004; Bach, 2015)*. To construct a map of intercellular signaling among heart cells using scRNA-seq data, we integrated them with a ligand-receptor interaction database (Ramilowski et al., 2015). Results showed that the endothelial and fibroblast clusters are prominent hubs for autocrine and paracrine signaling (Figure 2C and D, Supplementary file 9, Supplementary file 10, FDR≤0.01, log2FC ≥1), and that intercellular signaling in response to HFD/STZ-induced diabetes was changed in all cardiac cells, with fibroblasts increasing the largest number of connections (Figure 2E-G, Supplementary file 11, Supplementary file 12, 2-sided Wilcoxon rank-sum test, FDR≤ 0.05, log2FC≥ 0.36). Together, these analyses suggested that intercellular communications play an important role of the alterations of cardiac microenvironment of diabetic mice.

### Identification of key ligand-receptor pairs associated with diabetic myocardial fibrosis in fibroblasts

Cardiac fibroblasts are the primary drivers of myocardial fibrosis *(Travers et al., 2016; Frangogiannis, 2021)*. Given the result of cardiac fibroblasts increasing the greatest number of connections in response to HFD/STZ-induced diabetes, perturbations of intercellular communications between cardiac fibroblasts and cardiac populations may be key drivers of diabetic myocardial fibrosis. To investigate the key receptor-ligand interactions in diabetic myocardial fibrosis, the highly expressed receptors in cardiac fibroblasts were screened (Figure 3-Figure supplement 1A, FDR≤0.01, log2FC≥1).

**Figure 3.**
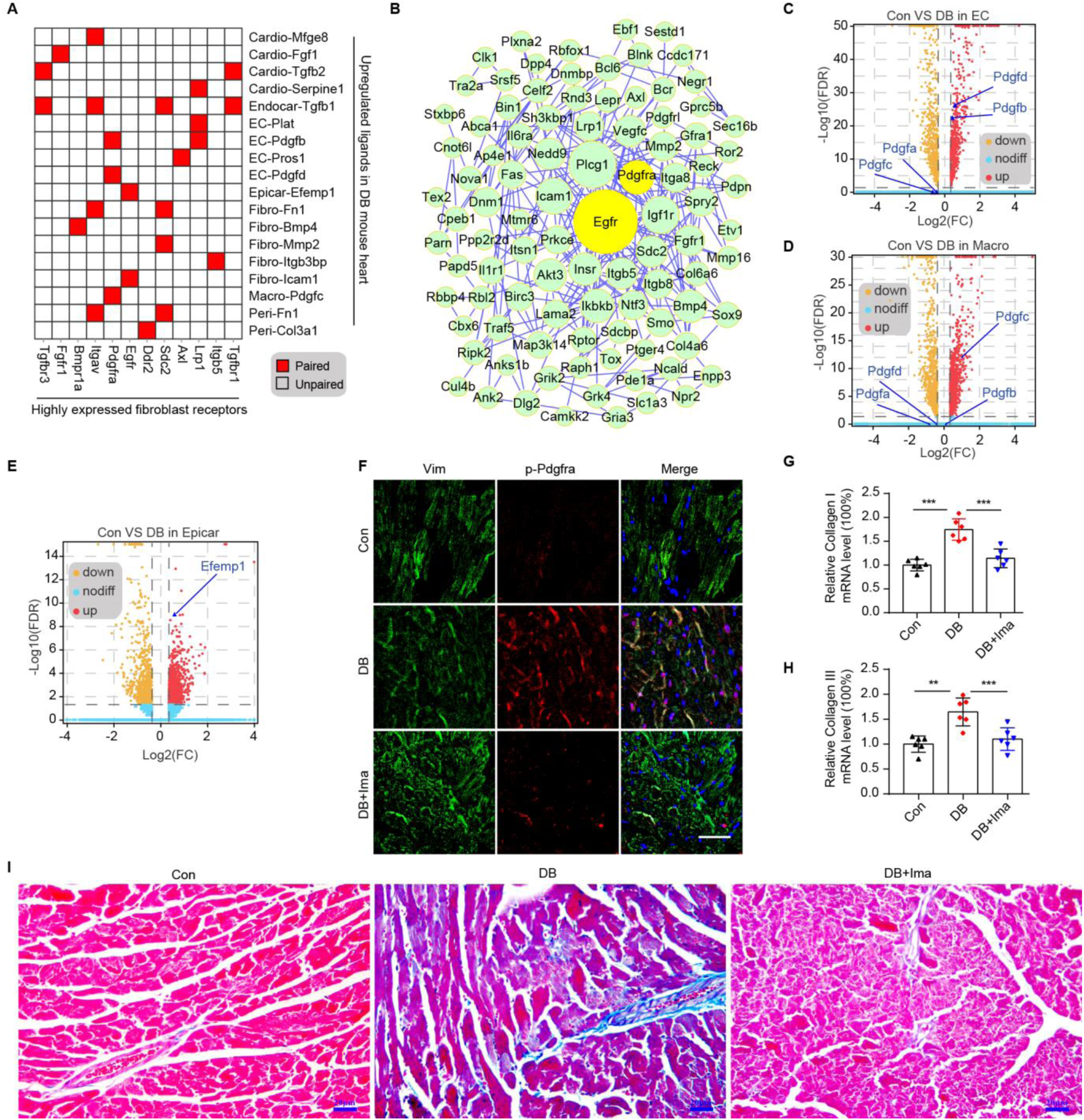
Identification of key ligand-receptor pairs associated with diabetic myocardial fibrosis in fibroblasts. (A) Heatmap shows pairs of highly expressed fibroblast receptors and the upregulated ligands in each cell type of diabetic hearts. (B) PPI network shows interaction of up-regulated genes in fibroblasts. The circle size represents the protein node degree in the network. (C, D) Volcano plots of DEGs in the heart tissues of HFD/STZ-treated and control mice. Pdgfa, Pdgfb, Pdgfc and Pdgfd in endothelial cells (C) and macrophages (D) are highlighted (2-sided Wilcoxon rank-sum test, FDR≤0.05, log2FC≥0.36). (E) Volcano plots of DEGs in the hearts from HFD/STZ-treated and control mice. Efemp1 in epicardial cells is highlighted (2-sided Wilcoxon rank-sum test, FDR≤0.05, log2FC≥0.36). (F) Representative immunofluorescence images for p-Pdgfra in heart tissues from HFD/STZ-treated mice with or without Ima treatment (n = 6 mice per group), scale bar = 40 µm. (G, H) Bar plots show mRNA expression of Collagen I (G) and collagen III (H) in heart from HFD/STZ-treated mice with or without Ima treatment. (n = 6 mice per group; mean ± SEM, **p < 0.01, ***p < 0.001). (I) Representative images of Masson dye-stained heart sections from the indicated groups showing extent of collagen deposition, (n = 6 mice per group), scale bar = 20µm. Ima, imatinib mesylate; SEM, standard Error of Mean. The details of cognate ligands of Egfr and Pdgfra are listed in *Supplementary file 13* and *Supplementary file 14*, respectively. This paper includes the following source data and figure supplement(s) for Figure 3. **Source data 1.** Source data for CT values of Collagen I used for Figure 3G. **Source data 2.** Source data for CT values of Collagen III used for Figure 3H. **Figure supplement 1.** Identification of key ligand-receptor pairs associated with diabetic myocardial fibrosis in fibroblasts. **Figure supplement 2.** The cognate ligands of Egfr in each cardiac cell population. **Figure supplement 3.** Immunofluorescence results of Pdgfb, Pdgfc, and Pdgfd in Pdgfra^+^ cells.

**Figure 3-Figure supplement 1.**
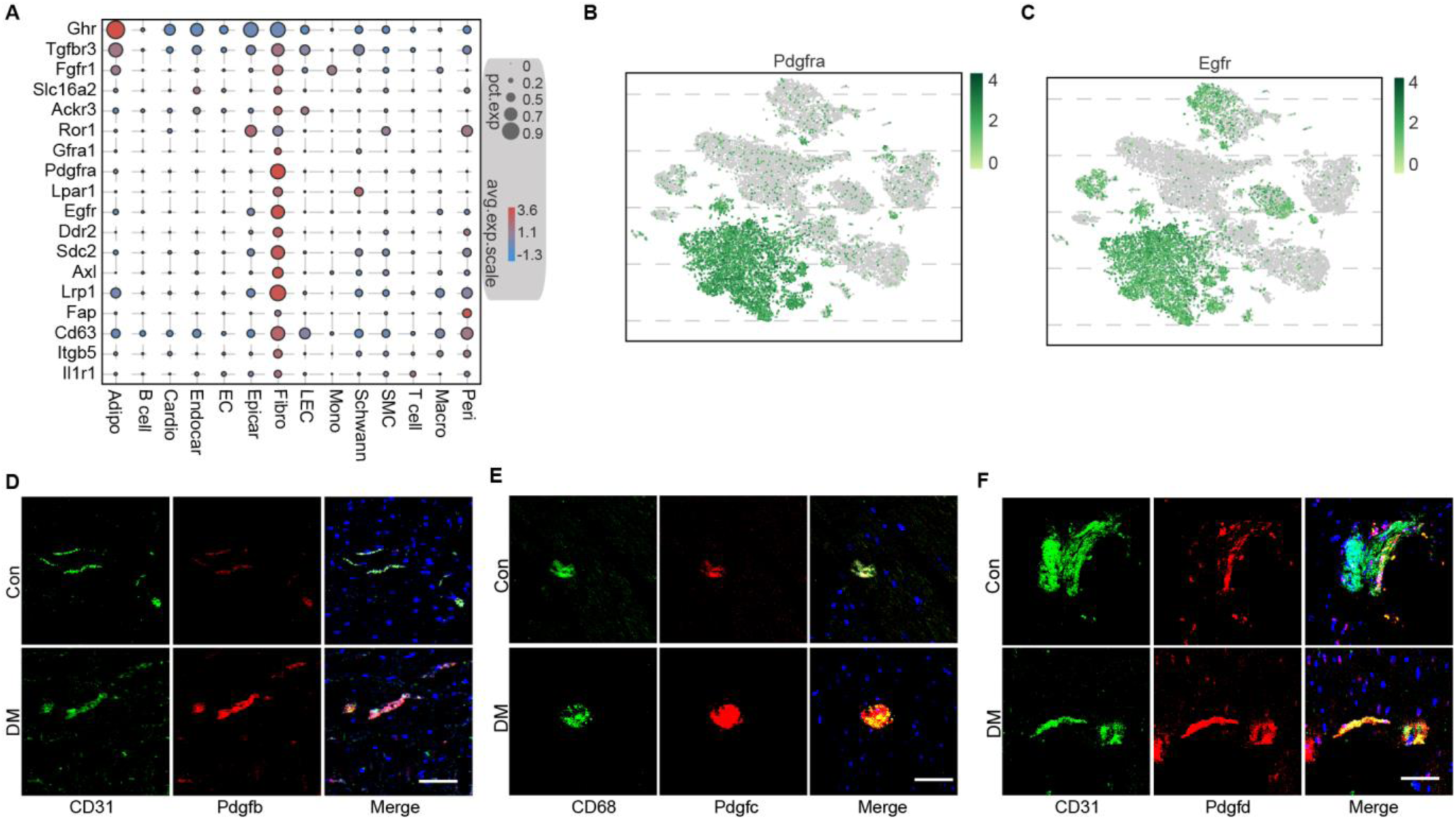
Identification of key ligand-receptor pairs associated with diabetic myocardial fibrosis in fibroblasts. (A) Dot plot shows the highly expression of specific receptors in cardiac cell populations (FDR≤0.01, log2FC≥1). The circle size indicates the proportion of cells within groups that express each transcript. The red and blue dots respectively indicate high and low expressed genes. (B, C) 2-dimensional t-SNE projection of Pdgfra (B) and Egfr (C) expression in cardiac cell populations. Green and grey colors respectively indicate the high and low expressed genes. (D-F) Representative immunofluorescence images for Pdgfb (D) and Pdgfc (E) in CD31+ cells, and Pdgfd (F) in CD68+ cells in the heart tissues of HFD/STZ-treated and control mice (n = 6 mice per group), scale bar = 100 µm.

**Figure 3-Figure supplement 2.**
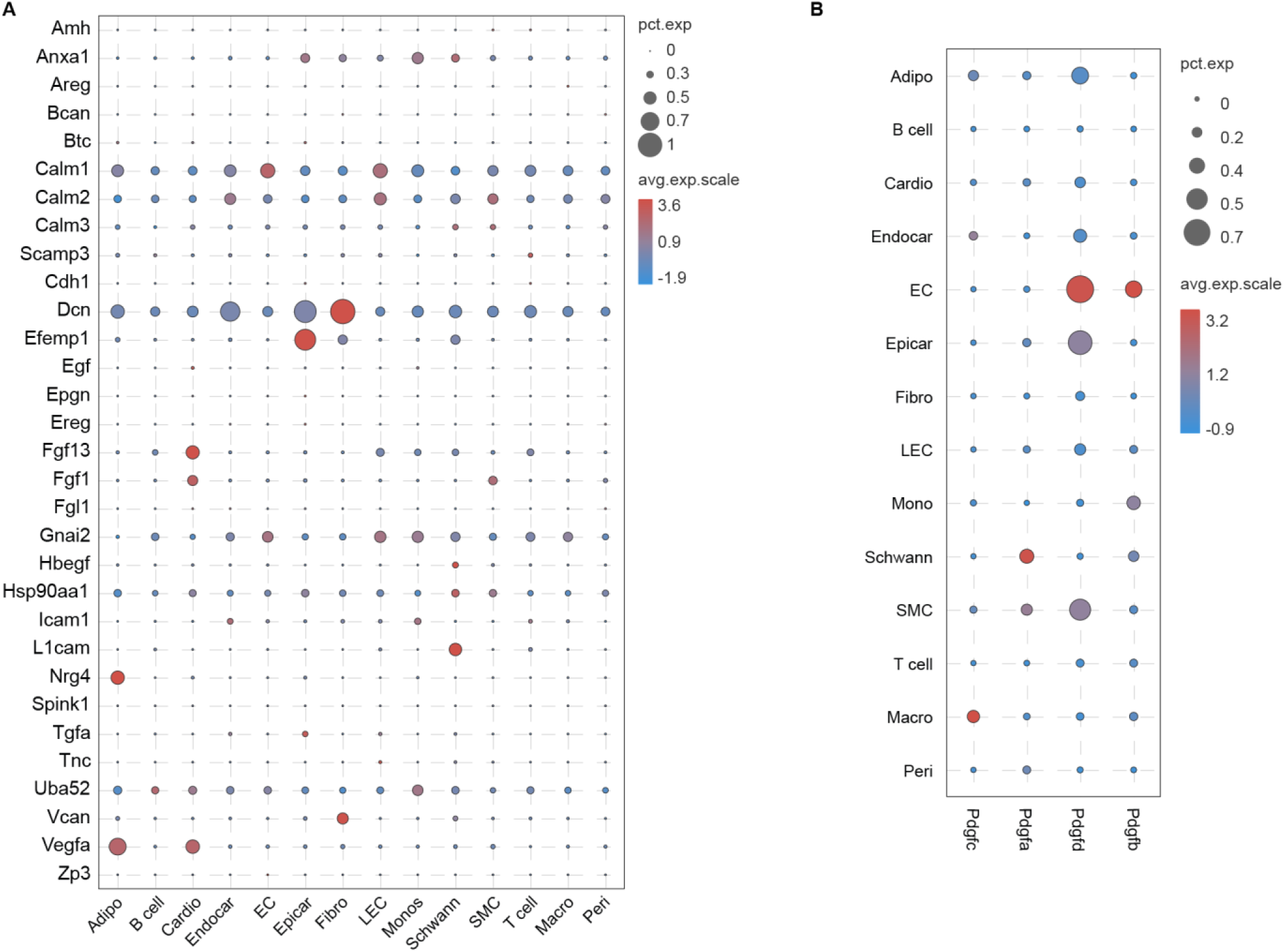
The cognate ligands of Egfr in each cardiac cell population. (A) Dot plot showing the cognate ligands of Egfr in each cardiac cell population. (B) Dot plot shows the cognate ligands of Pdgfra in cardiac cell populations. The circle size indicates the proportion of cells within groups that express each transcript. The red and blue dots respectively indicate high and low expressed genes.

**Figure 3-Figure supplement 3.**
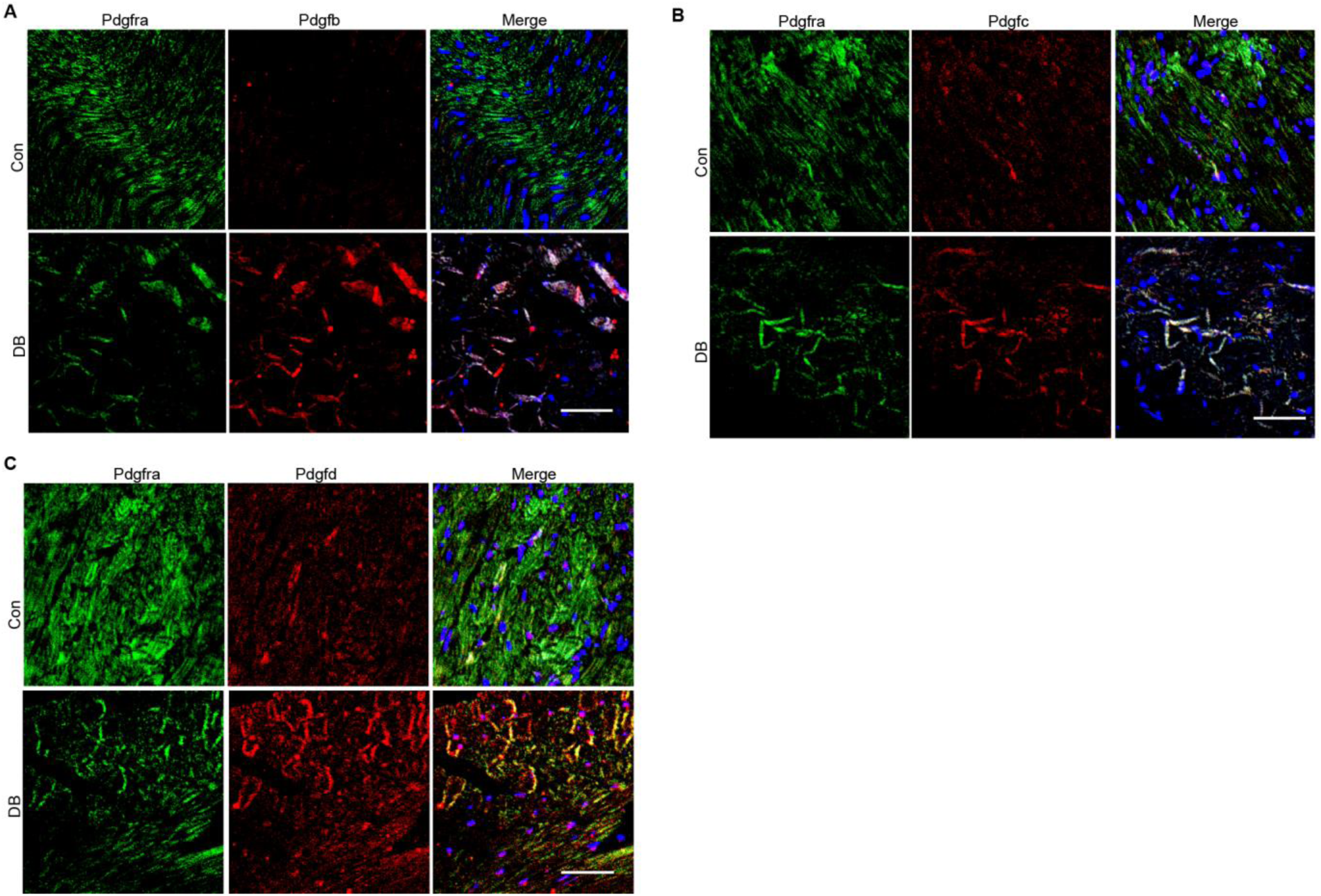
Immunofluorescence results of Pdgfb, Pdgfc, and Pdgfd in Pdgfra^+^ cells. (A-C) Representative immunofluorescence images for Pdgfb (A), Pdgfc (B) and Pdgfd (C) in Pdgfra^+^ cells in the heart tissues of HFD/STZ-treated and control groups (n = 6 mice per group), scale bar = 40 µm.

And then, we merged the upregulated uni-DEGs in fibroblasts and the highly expressed fibroblast receptors, whose cognate ligands were upregulated in at least one cardiac cell type during diabetic progression (Figure 3A). A protein-protein interaction (PPI) network was constructed using the new fibroblast-specific gene set (Figure 3B). Among the top hub genes based on the node degree were Egfr and Pdgfra, which were specifically high-expressed receptors in the cardiac fibroblasts (Figure 3-Figure supplement 1B and C). To clarify their role in fibrotic progression, we screened for the cognate ligands of Egfr and Pdgfra in each cardiac cell population (Figure 3-Figure 2A and B, Supplementary file 13 and Supplementary file 14). Both Pdgfb and Pdgfd transcripts were upregulated in endothelial cells, and Pdgfc levels was markedly increased in the macrophages of diabetic mice (Figure 3C and D, 2-sided Wilcoxon rank-sum test, FDR≤0.05, log2FC≥0.36). The protein levels of Pdgfb, Pdgfd, and Pdgfc showed similar changes in the corresponding cardiac cells (Figure 3-Figure supplement 1D-F, n = 6 mice per group). In addition, the Egfr ligand Efemp1 was upregulated in epicardial cells (Figure 3E, 2-sided Wilcoxon rank-sum test, FDR≤0.05, log2FC≥0.36). These results suggest that the interaction of cardiac fibroblasts with endothelial cells, macrophages and epicardial cells through Pdgf(s)-Pdgfra and Efemp1-Egfr may contribute to myocardial fibrosis in diabetic mouse heart.

Pdgfra exerts its tyrosine kinase activity through binding with its cognate ligands.

Immunostaining of the cardiac tissues revealed significantly higher protein levels of Pdgfb, Pdgfc and Pdgfd in the Pdgfra positive cells of the diabetic group (Figure 3-Figure supplement 3A-C, n = 6 mice per group). To examine the functional role of Pdgfra in diabetic myocardial fibrosis, we treated HFD/STZ-induced diabetic mice with Pdgfra inhibitor imatinib mesylate (Ima). Results showed that HFD/STZ treatment significantly increased cardiac p-Pdgfra protein levels and decreased that of p-Pdgfra in the HFD/STZ + Ima group compared to the HFD/STZ group (Figure 3F, n = 6 mice per group). In addition, the myocardium of the HFD/STZ-treated mice expressed higher levels of collagen I and III compared with the control group (Figure 3G and H, n = 6 mice per group, mean ± SEM, **p < 0.01, ***p < 0.001), which coincided with increased collagen deposition in the extracellular matrix. However, Ima treatment attenuated the increase in HFD/STZ-induced collagen synthesis and deposition (Figure 3I, n = 6 mice per group). Taken together, Pdgf(s)-Pdgfra interaction contributes to diabetic myocardial fibrosis.

### Identification of myocardial fibrosis-related cardiac fibroblast subpopulation

Cell subpopulations in tissues have non-overlapping functions and play different biological roles. *(Croft et al., 2019)*. Cell types can be defined by unbiassed clustering of single cells based on the global transcriptome patterns *(Rozenblatt-Rosen et al., 2017; McLellan et al., 2020)*. To observe heterogeneity of fibroblasts in heart, we examined 6416 fibroblasts from the diabetes and control mice. Unsupervised Seurat-based clustering of the 6416 fibroblasts revealed ten distinct subpopulations (Figure 4A, Supplement file 15, n = 3428 fibroblasts from healthy control and n = 2988 fibroblasts from 6 diabetic mice). Next, hierarchical clustering with multiscale bootstrap resampling was used to analyze the heterogeneity of these cardiac fibroblast subpopulations. This analysis revealed that fibroblast subpopulation 3 formed a distinct cluster based on its expression pattern from other fibroblast subpopulations (Figure 4B).

**Figure 4.**
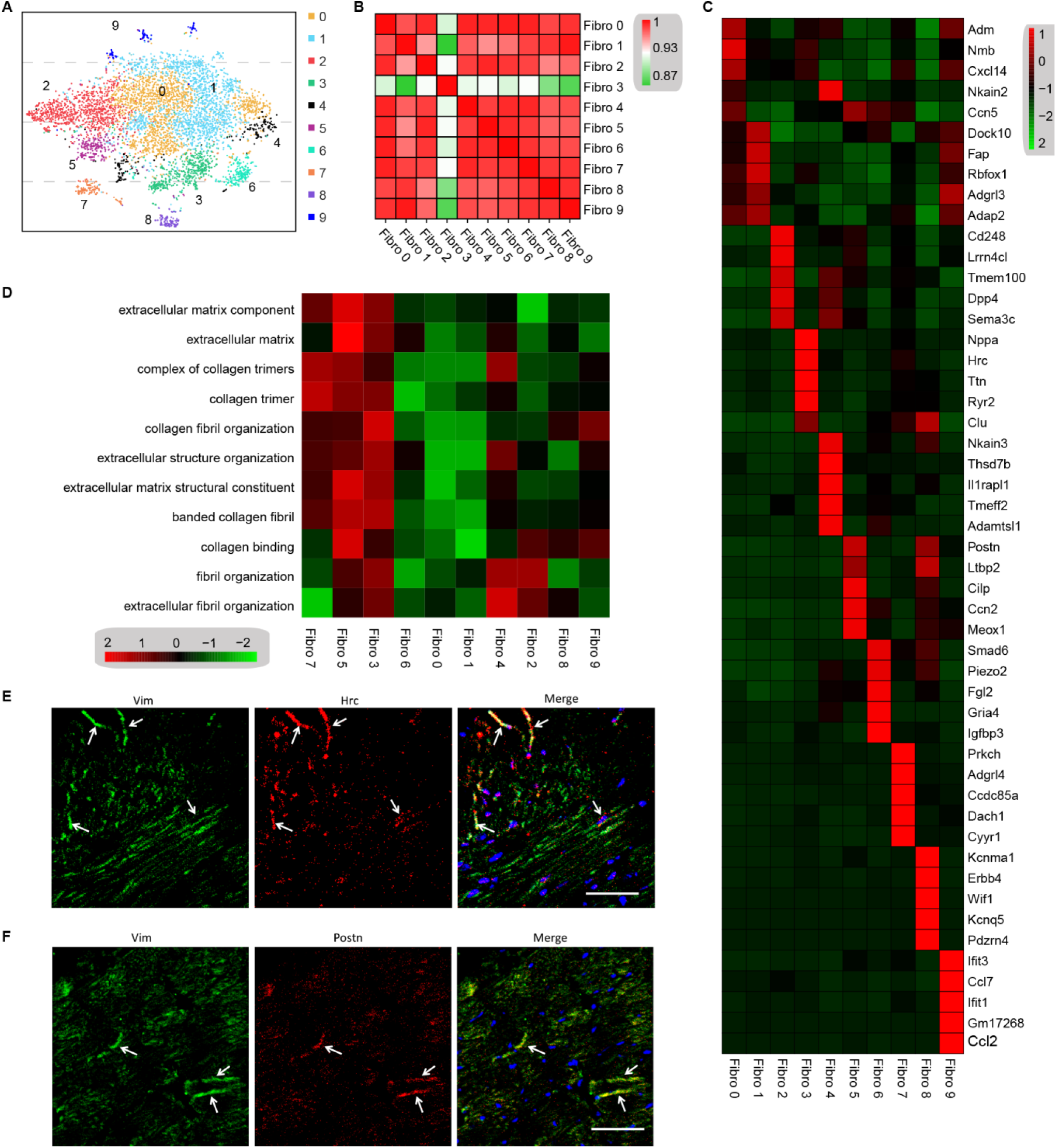
Analysis of the heterogeneity of fibroblast subpopulations. (A) t-SNE plot of ten cardiac fibroblast subpopulations from HFD/STZ-treated and control mice (n = 3428 fibroblasts from healthy control, and n = 2988 fibroblasts from 6 diabetic mice). (B) Correlation heatmap of gene-expression signatures of each fibroblast subpopulation. Color differences indicate subpopulations that were resolved by multiscale bootstrapping. (C) Heatmap shows the top five marker genes for each fibroblast subpopulation (FDR≤ 0.05, log2FC ≥ 0.36). Red color indicates high expression; green color indicates low expression. (D) Heatmap shows the enriched GO terms associated cardiac fibrosis in each fibroblast population (FDR≤0.05). Color scale: red, low FDR; green, high FDR. (E, F) Representative immunofluorescence images for Hrc (F) and Postn (G) in mouse heart (n = 6 mice per group), scale bar = 100 µm. The details of 10 transcriptionally distinct fibroblast subpopulations are listed in *Supplementary file 15*. The details of distinct signatures of each fibroblast subpopulations in heart are listed in *Supplementary file 16*.

We further investigated the contribution of all fibroblast populations to myocardial fibrosis. The top-5 ranking markers from the heart showed distinct signatures for each subpopulation of fibroblasts by heatmap analysis (Figure 4C, Supplementary file 16, FDR≤0.05, log2FC≥0.36). Of note, the top enriched genes in subpopulation 3 (Nppa and Clu) and subpopulation 5 (Postn and Cilp) are well-established biomarkers of pro-fibrotic function. Gene set variation analysis (GSVA) of each fibroblast subpopulation suggested a diversification of function between the subpopulations, and fibroblast 3 and 5 populations were strongly involved in extracellular matrix remodeling and collagen synthesis (Figure 4D, FDR≤0.05). These results indicated that fibroblast 3 and 5 subpopulations are myocardial fibrosis-related cardiac fibroblast subpopulations.

The most significantly enriched gene in subpopulation 3 was Hrc, which is crucial for proper cardiac function by regulating Ca^2+^-uptake, storage and release. The most significantly enriched genes of fibroblast subpopulation 5 was Postn, which is consistent with a fibroblast subset identified in an animal model of angiotensin-induced myocardial hypertrophy *(McLellan et al., 2020)*. The Hrc^hi^ and Postn^hi^ fibroblast populations were also detected in mouse heart by immunostaining for Hrc and Postn respectively (Figure 4E and F, n = 6 mice per group).

### Transcription Factor Network Analysis

To investigate the underlying molecular mechanisms that drive the phenotypic differentiation of fibroblast subpopulations, we used single-cell regulatory network inference and clustering. Results revealed upregulation of different transcription factor networks in the distinct subpopulations. For instance, Thra and Creb5 regulons were upregulated in fibroblast subpopulation 0 and 2 respectively, whereas the Nfe2l1 network was enriched in subpopulation 3 and subpopulation 4 showed increased activation of the Foxp2 network (Figure 5A, Supplementary file 17). Regulons driven by the Tcf4 transcription factors were enriched in subpopulations 1 and 9, and Mef2a was enriched in subpopulations 7 and 8. Consistent with the role of Hmgb1 in cardiac fibrosis *(Wu et al., 2018)*, a Hmgb1-based network was upregulated in Hrc^hi^ and Postn^hi^ fibroblast populations (Figure 5A). Heatmap analysis further confirmed these upregulated transcription factors (Figure 5B, Supplementary file 18).

**Figure 5.**
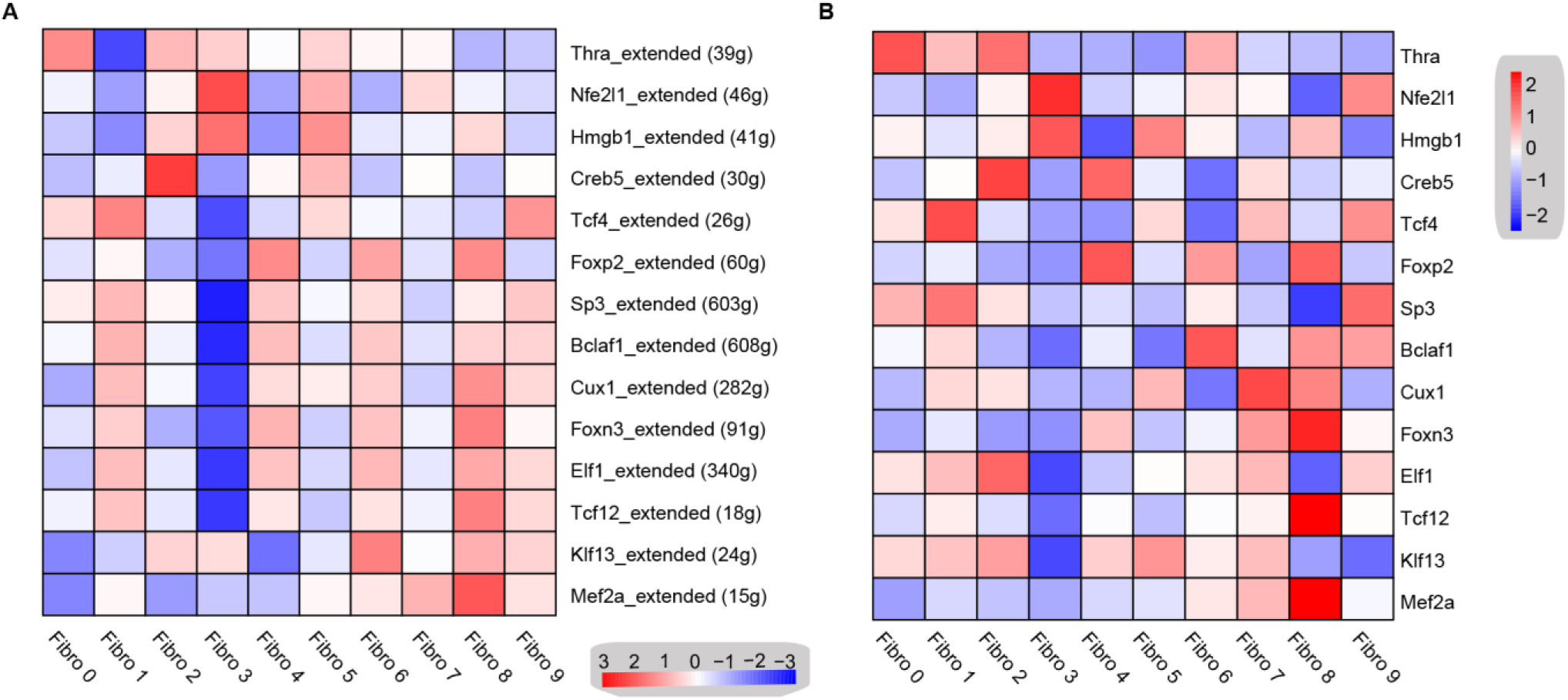
Transcription factor network analysis of fibroblast subpopulations. (A) Heatmap shows the inferred transcription-factor gene-regulatory networks. Numbers between brackets indicate the (extended) regulons for respective transcription factors. (B) Heatmap shows the expression level of transcription factors in (A). Red color indicates high expression; green color indicates low expression. The details of transcription-factor gene-regulatory networks in the distinct subpopulations are listed in *Supplementary file 17*. The details of transcription factors expression are listed in *Supplementary file 18*.

### Identification of intercellular communication drivers of myocardial fibrosis in Hrc^hi^ fibroblasts

To identify the cellular drivers of myocardial fibrosis during diabetes, we performed a cluster analysis of the DEGs between control and diabetic mice heart. HFD/STZ treatment induced transcriptional changes in all cardiac fibroblast subpopulations (Figure 6-Figure supplement 1, Supplementary file 19, 2-sided Wilcoxon rank-sum test, FDR≤0.05, log2FC≥0.36), and the upregulated genes in Hrc^hi^ fibroblasts were enriched in GO terms such as collagen fiber reorganization and extracellular matrix binding (Figure 6A, 2-sided Wilcoxon rank-sum test, FDR≤0.05). Furthermore, the top 20 enriched pathways in the Hrc^hi^ fibroblasts of the diabetic group were related to extracellular matrix organization, myocardial fibrosis and fibroblast activation (Figure 6B, 2-sided Wilcoxon rank-sum test, FDR≤0.05). These results suggested that the Hrc^hi^ fibroblasts are the key cellular driver of myocardial fibrosis in response to HFD/STZ-induced diabetes.

**Figure 6.**
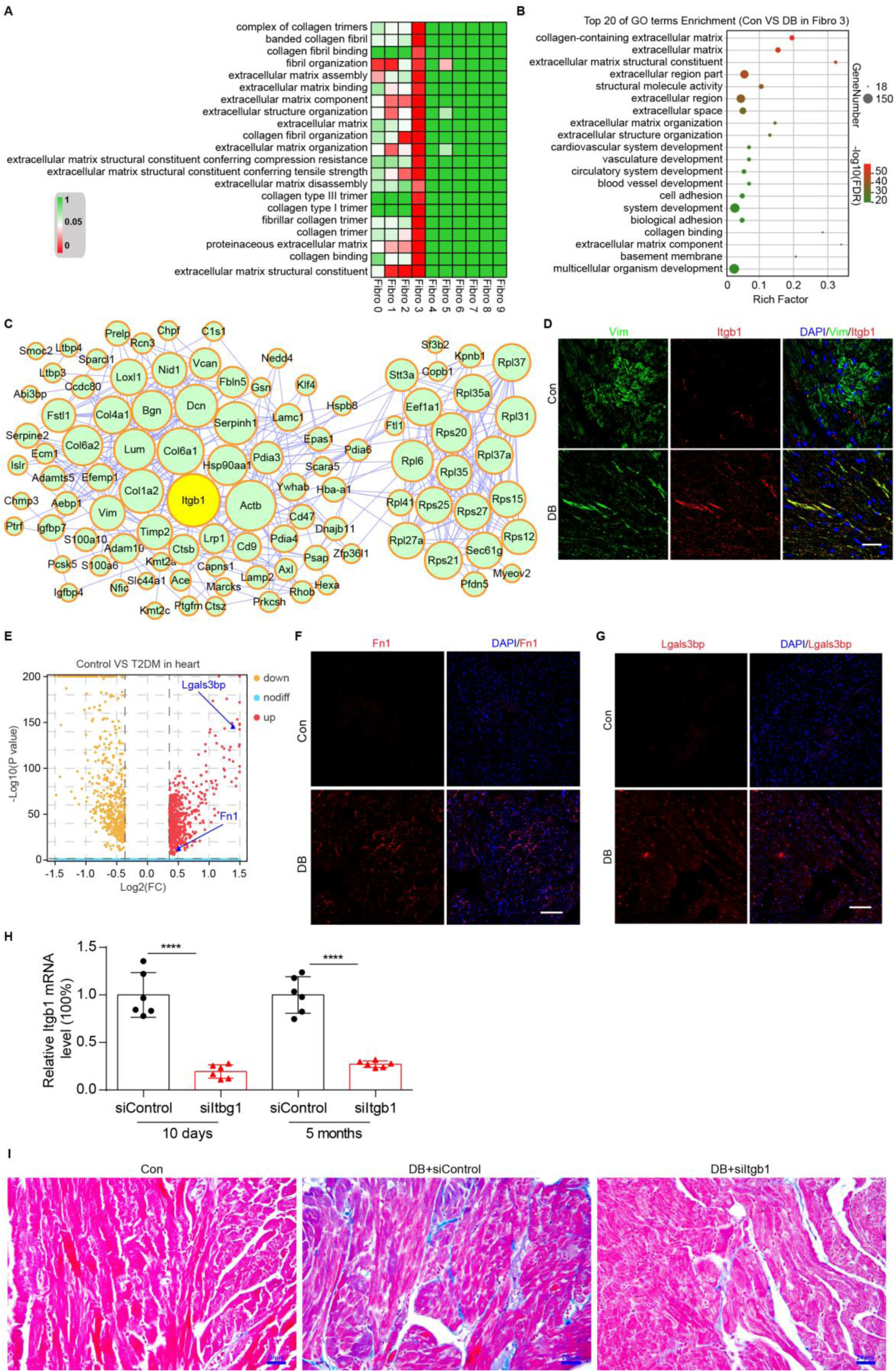
Identification of intercellular communication drivers of myocardial fibrosis in Hrc^hi^ fibroblasts. (A) Heatmap shows HFD/STZ-induced enrichment of GO terms associated with extracellular matrix remodeling and myocardial fibrosis in each subpopulation of cardiac fibroblasts (2-sided Wilcoxon rank-sum test, FDR≤0.05). Color scale: red, low FDR value; green, high FDR value. (B) Dot plot of GO analysis shows the top 20 terms with the highest enrichment in Hrc^hi^ fibroblasts in the HFD/STZ-treated mice relative to controls (2-sided Wilcoxon rank-sum test, FDR≤0.05). Bars are color-coded from blue to red based on the FDR. (C) PPI network shows interaction of up-regulated genes in Hrc^hi^ fibroblasts of diabetic mice relative to controls. The circle size represents the protein node degree in the network. (D) Representative immunofluorescence images for Itgb1 in heart from SHH-fed or control mice (n = 6 mice per group), scale bar = 40 µm. (E) Volcano plots shows DEGs in the heart tissues from HFD/STZ-treated or control mice with Fn1 and Lgals3bp highlighted (2-sided Wilcoxon rank-sum test, FDR≤0.05, log2FC≥0.36). (F, G) Representative immunofluorescence images for Fn1 (F) and Lgals3bp (G) in mouse heart (n = 6 mice per group), scale bar = 100 µm. (H) The efficiency of siRNA-mediated Itgb1 mRNA knockdown was confirmed by qRT–PCR (n = 6 mice per group, mean ± SEM, ****p < 0.0001). (I) Representative images of Masson dye-stained heart sections from the indicated groups showing extent of collagen deposition (n = 6 mice per group), scale bar = 20µm. Detailed genes of significant transcriptomic changes in each fibroblast subpopulation are listed in *Supplementary file 19*. The details of unique differentially-expressed genes (uni-DEGs) in each fibroblast subpopulation are listed in *Supplementary file 20*. The details of the cognate ligands of Itgb1 are listed in *Supplementary file 21*. This paper includes the following source data and figure supplement(s) for Figure 6. **Source data 1.** Source data for CT values of Itgb1 used for Figure 6H. **Source data 2.** Source data for CT values of Itgb1 used for Figure 6H. **Figure supplement 1.** Up- and downregulated genes in each fibroblast subpopulation of diabetic versus control mice. **Figure supplement 2.** DEGs in each cardiac fibroblast subpopulation from HFD/STZ-treated or control mice with Itgb1 highlighted.

**Figure 6-Figure supplement 1.**
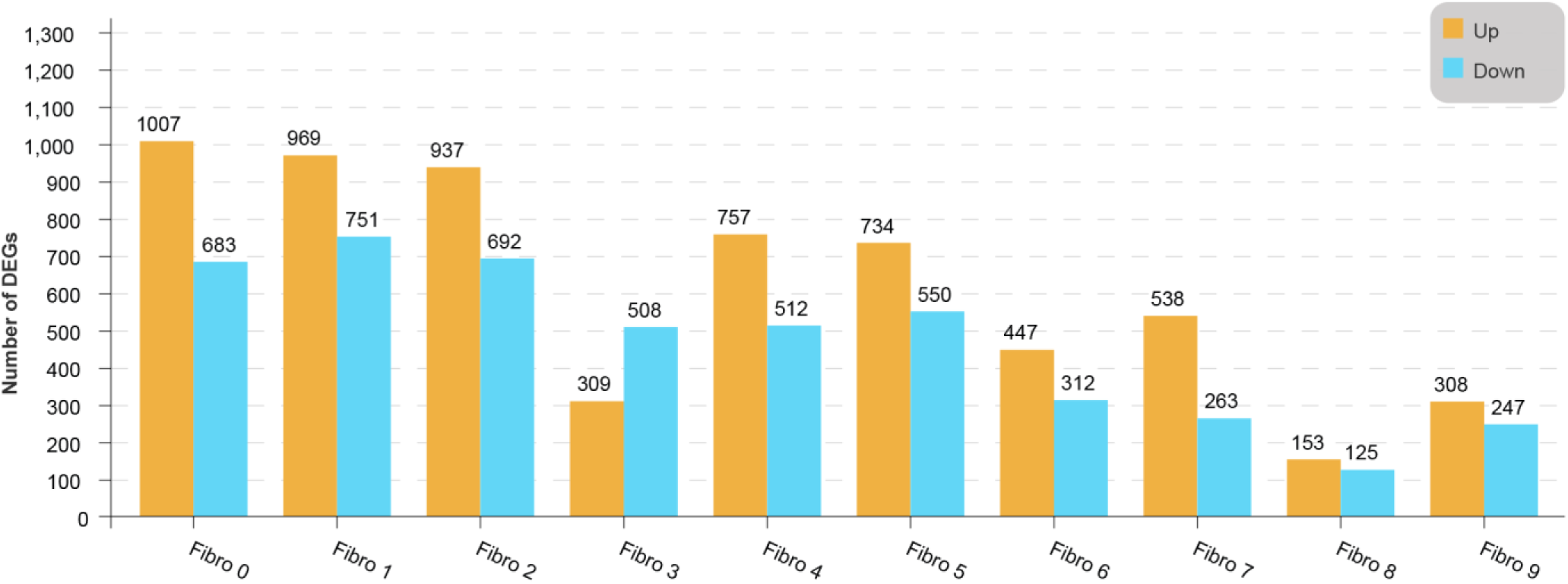
Lollipop plot shows the up- and downregulated genes in each fibroblast subpopulation of diabetic versus control mice (2-sided Wilcoxon rank-sum test, FDR≤0.05, log2FC≥0.36).

**Figure 6-Figure supplement 2.**
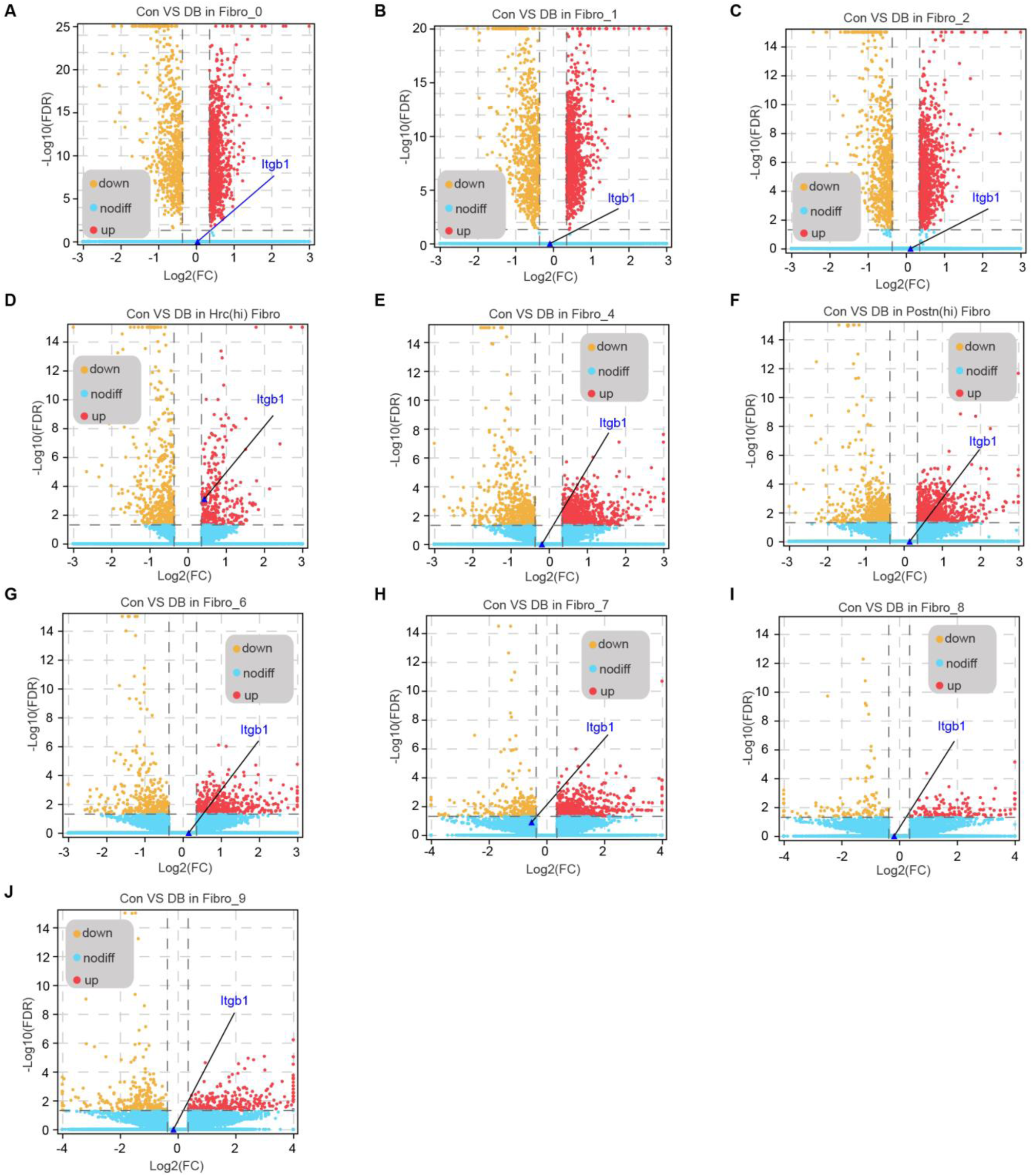
Volcano plots shows DEGs in each cardiac fibroblast subpopulation from HFD/STZ-treated or control mice with Itgb1 highlighted (2-sided Wilcoxon rank-sum test, FDR≤0.05, log2FC≥0.36).

To identify the critical signaling molecules in Hrc^hi^ fibroblasts that mediate myocardial fibrosis in the diabetic setting, we identified the uni-DEGs in each fibroblast subpopulation (Supplementary file 20) and constructed a PPI network using the upregulated genes (Figure 6C). The top 15 hub genes included Itgb1, Col6a1, Col1a2, Dcn, Rpl6, Rps20, Serpinh1, Bgn, Hsp90aa1 and Col6a2, of which Col6a1, Col1a2, Col6a2, Dcn and Bgn encode for ECM proteins *(Schipke et al., 2017)*. In addition, both Serpinh1 and Hsp90aa1 have been reported to participate in collagen synthesis *(Christiansen et al., 2010; García et al., 2016)*. Although the role of these candidate hub genes has been well established in myocardial fibrosis, the functions of Itgb1 is currently unknown.

The PPI network of the upregulated uni-DEGs indicated the key role of Itbg1 of Hrc^hi^ fibroblasts in diabetic myocardial fibrosis (Figure 6D, n = 6 mice per group; Figure 6E, Figure 6-Figure supplement 2, 2-sided Wilcoxon rank-sum test, FDR≤0.05, log2FC≥0.36). Itgb1 deficiency increased the risk of ventricular arrhythmias in patients with arrhythmogenic right ventricular cardiomyopathy (Wang et al., 2020). It is therefore reasonable to surmise that the interaction of Itgb1 with its cognate ligand is involved in diabetes-related myocardial fibrosis. To confirm this hypothesis, we screened all potential ligands of Itgb1 (Supplementary file 21), and found that Lgals3bp and Fn1 were upregulated in the heart tissues of diabetic mice (Figure 6F-G, n = 6 mice per group).

To validate the role of Itgb1 in heart for myocardial fibrosis during diabetes, we used the adeno-associated virus 9 (AAV9) to deliver Itgb1 siRNA, which preferentially target heart. The level of Itgb1 mRNA decreased by >80% in response to a single injection of Itgb1 siRNA compared to negative control (Figure 6H, n = 6 mice per group, mean ± SEM, ****p < 0.0001). Moreover, the knockdown of Itgb1 lasted for more than 5 months after injection of siRNA. Next, we tested the collagen synthesis and deposition in diabetic mouse heart with Itgb1 knockdown by Masson dye-staining. Results showed that levels of the collagen synthesis and deposition were indeed reduced in Itgb1 knockout mice (Figure 6I, n = 6 mice per group). Taken together, Itgb1 in Hrc^hi^ fibroblasts contributes to HFD/STZ-induced myocardial fibrosis. Further studies are warranted to establish the roles of these ligand-receptor interactions in diabetic myocardial fibrosis.

### Identification of SNPs of Itgb1 and Fn1 correlated with type 2 diabetes and glucose metabolic disorders

To investigate the clinical relevance of Itgb1 to type 2 diabetes and glucose metabolic disorders, we examined the gene polymorphism of Itgb1 in human subjects. We searched single nucleotide polymorphism (SNP) located in Itgb1 and its ligands, Fn1 and Lgals3bp, in GWAS CENTRAL (https://www.gwascentral.org/) and found dozens of SNPs correlated with type 2 diabetes and blood glucose parameters. As summarized in table 1, three SNPs, rs2230394, rs2230395, and rs16933819 located on Itgb1 are correlated with type 2 diabetes. Among them rs2230395 causes termination of translation (Tyr153=Y(Tyr)> *(Ter)) and rs2230395 results in synonymous variant of protein. Four SNP loci of Fn1 are correlated with type 2 diabetes, among them rs13652 causes termination of translation (1575Asp=E(Glu)>D(Asp)). Although other SNPs of Itgb1 and Fn1 correlated with type II diabetes cause intron variant and do not affect amino acids, they all have significant correlation with type 2 diabetes (P<0.05). There are no Lgals3ps SNP loci significantly correlated with type 2 diabetes (Table 1).

**Table 1.** SNPs of Itgb1 and Fn1 correlated with type 2 diabetes and glucose metabolic disorders.

| Correlations with type II diabetes |  |  |  |  |  |  |  |
| --- | --- | --- | --- | --- | --- | --- | --- |
| SNP | Gene | p-value | Alleles | Position | Consequence | Amino acid | Dataset identifier |
| rs2230394 | Itgb1 | 0.02823 | G>A,C | chr10:32928182 (GRCh38) | Stop Gained | NP_002202.2:p. Tyr153Ter<br>Y (Tyr) > *(Ter) | HGVRs9 |
| rs2230395 | Itgb1 | 0.03615 | T>A,G | chr10:32922299 (GRCh38) | Synonymous Variant | NP_002202.2:p. Ala362=<br>A(Ala) >A (Ala) | HGVRs9 |
| rs16933819 | Itgb1 | 0.0498 | T>C | chr10:32922299 (GRCh38) | Intron variant |  | HGVRs9 |
| rs13652 | Fn1 | 0.01 | C>A,T | chr2:215384864 (GRCh38) | Missense Variant | NP_997647.2:p. Glu1575Asp<br>E(Glu) > D(Asp) | HGVRs16 |
| rs1968510 | Fn1 | 0.047 | A>C,G, T | chr2:215393528 (GRCh38) | Intron variant |  | HGVRs16 |
| rs2372544 | Fn1 | 0.031 | G>A,C, T | chr2:215399547 (GRCh38) | Intron variant |  | HGVRs16 |
| rs6744921 | Fn1 | 0.036 | A>G | chr2:215402013 (GRCh38) | Intron variant |  | HGVRs16 |
| Correlations with fasting insulin-related: fasting insulin |  |  |  |  |  |  |  |
| SNP | Gene | p-value | Alleles | Position | Consequence | Amino acid | Dataset identifier |
| rs2245844 | Itgb1 | 0.04343 | C>A,T | chr10:32935848 (GRCh38.p13) | Intron Variant |  | HGVRs3266 |
| rs2488329 | Itgb1 | 0.04549 | A>C,G, T | chr10:32926545 (GRCh38.p13) | Intron Variant |  | HGVRs3266 |
| rs2503995 | Itgb1 | 0.04766 | G>A,C | chr10:32924543 (GRCh38.p13) | Intron Variant |  | HGVRs3266 |
| rs2503996 | Itgb1 | 0.04557 | C>T | chr10:32924909 (GRCh38.p13) | Intron Variant |  | HGVRs3266 |
| Correlations with fasting glucose-related: fasting plasma glucose |  |  |  |  |  |  |  |
| rs11883812 | Fn1 | 0.03278 | T>C | chr2:215392258<br>(GRCh38.p13) | Intron<br>Variant |  | HGVRs3269 |
| rs17516906 | Fn1 | 0.0473 | A>G | chr2:215414550<br>(GRCh38.p13) | Intron<br>Variant |  | HGVRs3269 |
| rs17517928 | Fn1 | 0.03306 | C>T | chr2:215426636 | Intron<br>Variant |  | HGVRs3269 |
| **Correlations with Fasting glucose-related: homeostatic model assessment of beta-cell function** |  |  |  |  |  |  |  |
| rs1250201 | Fn1 | 0.0452 | C>A,G,<br>T | chr2:215385096<br>(GRCh38) | Intron Variant |  | HGVRs3267 |
| rs1250203 | Fn1 | 0.01508 | G>C,T | chr2:215383582<br>(GRCh38.p13) | Intron Variant |  | HGVRs3267 |
| rs13652 | Fn1 | 0.01951 | C>A,T | chr2:215384864 | Missense<br>Variant | NP_997647.2:p.<br>Glu1575Asp<br>E (Glu) >D(Asp) | HGVRs3267 |
| rs1968509 | Fn1 | 0.03908 | C>G,T | chr2:215390542<br>(GRCh38.p13) | Intron Variant |  | HGVRs3267 |
| rs1968510 | Fn1 | 0.007892 | A>C,G,<br>T | chr2:215393528<br>(GRCh38.p13) | Intron Variant |  | HGVRs3267 |
| rs2577302 | Fn1 | 0.0278 | T>A,C,<br>G | chr2:215392542<br>(GRCh38.p13) | Intron Variant |  | HGVRs3267 |
| rs2692230 | Fn1 | 0.0227 | C>A,G,<br>T | chr2:215392624<br>(GRCh38.p13) | Intron Variant |  | HGVRs3267 |
| rs6435904 | Fn1 | 0.04018 | T>A,C,<br>G | chr2:215389839<br>(GRCh38.p13) | Intron<br>Variant |  | HGVRs3267 |

SNPs of Itgb1 and Fn1 are also correlated with other blood glucose parameters. Eight SNP loci of Fn1 are associated with fasting glucose-related: homeostatic model assessment of beta-cell function, among them rs13652 causes missense variant of Fn1 (1575Asp=E(Glu)>D(Asp)) (P=0.01951). Four SNPs of Itgb1 are associated with fasting insulin-related: fasting insulin (P<0.05). Three SNPs of Fn1 are associated with fasting glucose-related: fasting plasma glucose (P<0.05) (Table 1). Together, these GWAS data supported the role of Fn1-Itgb1 pair in type 2 diabetes.

## Discussion

scRNA-seq allows in-depth analysis of the individual cells in heterogenous populations *(Butler et al., 2018)* of healthy and diseased tissues *(Mathys et al., 2019; Peng et al., 2019; Kalucka et al., 2020; Litviňuková et al., 2020; Li et al., 2021)*. However, the effect of diabetes on cardiac cell function and cardiac cell heterogeneity at single-cell level has not been reported so far. In this study, we mapped the transcriptional alterations associated with HFD/STZ-induced diabetes in different cardiac cell populations, and identified the key ligand-receptor pair drivers of myocardial fibrosis in fibroblasts of diabetic heart. Specifically, the emergence of Hrc^hi^ fibroblast subpopulations in response to diabetic progression, presumably to remodel the extracellular environment by multiple ligand-receptor interactions.

The heart of a mammal is a complex organ composed of a variety of cell types *(Banerjee et al., 2007; Litviňuková et al., 2020)*. Cardiac fibroblasts synthesize extracellular matrix proteins and their excessive activation in response to stress induces cardiac fibrosis *(Travers et al., 2016)*. GO enrichment analysis of all upregulated genes in cardiac cell populations from the diabetic mice confirmed the association between fibroblasts and extracellular matrix remodeling and myocardial fibrosis, which is consistent with previous studies *(Jia et al., 2018; Ivey et al., 2019; McLellan et al., 2020; Frangogiannis, 2021)*. The survival and proper functioning of metazoans depends on the communication between multiple cell populations and tissues via secretary ligands and membrane receptors *(Ramilowski et al., 2015)*. To this end, we screened for the receptor genes that were highly expressed in cardiac fibroblasts of diabetic mice and their cognate ligands that were upregulated in other cardiac cell populations to identify the dysregulated ligand-fibroblast receptor interactions. Protein-protein interaction network analysis indicated that the receptors Pdgfra and Egfr, which are highly expressed in fibroblasts, play central roles in myocardial fibrosis of diabetes. Pdgfra is a surface receptor tyrosine kinase that is activated upon binding to its corresponding ligand Pdgf(s), and regulates cell division and proliferation *(Rudat et al., 2013; Gouveia et al., 2018; Soliman et al., 2020)*. Egfr on the other hand is a member of the ErbB family of receptor tyrosine kinases and plays an important role in wound healing and cardiac hypertrophy *(Peng et al., 2016)*. The upregulation of Pdgfra ligands Pdgfb and Pdgfd in endothelial cells, and Pdgfc in macrophages, and that of the Egfr ligand Efemp1 in epicardial cells of the HFD/STZ-treated mice indicated that the cardiac microenvironment was changed, resulting in extracellular matrix remodeling and cardiac fibrosis. This is of particular interest given the pathological roles of these cell populations and ligand-receptor pairs in cardiovascular diseases *(Rottlaender et al., 2011; Shinagawa and Frantz, 2015; Farbehi et al., 2019; Marín -Juez et al., 2019; Peet et al., 2020; Baguma-Nibasheka et al., 2021)*. Ligand-receptor pair analysis also revealed a synergistic role of endothelial cells, macrophages and epicardial cells with the fibroblasts in diabetic myocardial fibrosis. Further analysis of the interactions between these cell populations will help understand the pathogenesis of diabetes-induced fibrosis.

Terminally differentiated cells are generally considered to have limited plasticity. Most cellular plasticity in adults is reported in the terminal differentiation stage of many progenitor cells *(Chang-Panesso and Humphreys, 2017)*. However, these cellular transitions also may be present in cardiac fibroblasts. Unbiased single-cell clustering can redefine cell types on basis of the global transcriptome patterns *(Rozenblatt-Rosen et al., 2017; McLellan et al., 2020)*. Such analyses have already been applied to other organs *(Macosko et al., 2015; Chen et al., 2017; Stubbington et al., 2017)* and even to whole multicellular organisms *(Cao et al., 2017; Karaiskos et al., 2017)*. These experiments have identified new cells as well as previously defined cells with catalogued marker genes, demonstrating that this approach has the ability to redefine cardiac cell types. One of the most important results of mouse cardiac fibroblasts analysis was the identification of two different phenotypic Hrc^hi^ and Postn^hi^ fibroblast subpopulations that were associated with extracellular matrix remodeling. The Postn^hi^ fibroblasts participate in fibrogenic progression, which is consistent with another subpopulation of fibroblasts identified in an animal model of angiotensin-induced myocardial hypertrophy *(McLellan et al., 2020)*, suggesting that these fibroblasts may contribute to both myocardial hypertrophy and cardiac fibrosis. Hrc^hi^ fibroblasts expressed fibrogenic marker genes such as Nppa, Ttn and Clu, which points to a key pro-fibrotic function. Hrc knockout or AAV mediated knockdown result in pulmonary edema, severe cardiac hypertrophy, fibrosis, heart failure and decreased survival after transverse aortic constriction (TAC) in mice *(Park et al., 2012, 2013)*. Combined with our single cell sequencing results, we can surmise that Hrc is a potential target for inhibiting myocardial fibrosis during diabetes.

Strikingly, GSVA and GO analyses of each fibroblast subpopulation indicated that Hrc^hi^ fibroblasts were the most profibrogenic under diabetic conditions. This finding suggests that the Hrc^hi^ fibroblasts may be the key cellular drivers of myocardial fibrosis in diabetes. We speculate that the intercellular communications between Hrc^hi^ fibroblasts and other cardiac cells are a constitutive process that is activated in diabetes. Intercellular and protein-protein interaction networks within the Hrc^hi^ fibroblasts reveal the key role of receptor Itgb1 in diabetic myocardial fibrosis. And the potential ligands of Itgb1, Lgals3bp and Fn1, are upregulated in the heart tissues of diabetic mice. Perhaps, the gain-of-function of Lgals3bp-Itgb1 and Fn1-Itgb1 pairs explains the role of Hrc^hi^ fibroblasts in diabetic myocardial fibrosis.

To explore the clinical significance of the role of Itgb1 in type 2 diabetes, we searched GWAS database and identified dozens of SNPs located on Itgb1 that are correlated with type 2 diabetes and other blood glucose parameters in human population. Although most SNPs identified are non-coding and only cause intron variant, they may regulate gene expression via modification of promoter and enhancer activity or disruption of binding sites for transcription factors *(Cano-Gamez and Trynka, 2020)*. Besides Itgb1, we also identified several SNPs located on Fn1 correlated with type 2 diabetes and other blood glucose parameters, while we only verified the role of Itgb1 in myocardial fibrosis of diabetes. The GWAS data supported the role of Itgb1 and Fn1 in type 2 diabetes in human population. Whole exon sequencing and other bioinformatic tools may help to identify additional amino acid variants of Itgb1 and its ligands in type 2 diabetes and related diseases.

In summary, we mapped the transcriptional alterations associated with HFD/STZ-induced diabetes in different cardiac subpopulations, and identified the key ligand-receptor pair drivers of myocardial fibrosis in diabetic heart, specifically the Pdgf(s)-Pdgfra and Efemp1-Egfr interaction mediated by fibroblasts with macrophages, endothelial cells and epicardial cells respectively. Crucially, Hrc^hi^ fibroblasts were identified as the key profibrogenic subpopulation that may contribute to cardiac fibrosis by remodeling the extracellular environment through the drivers of intercellular communication mediated by Itgb1. Therefore, we speculate that specific targeting Hrc^hi^ fibroblasts will be a promising target for the treatment of myocardial fibrosis. Our future research direction will entail combining fibroblast-specific Hrc knockout mice and analysis of the cardiac cellular networks to verify the role of Hrc in regulating diabetic myocardial fibrosis.

## Materials and methods

### Animals and treatments

Male C57BL/6J mice weighing 18-22 g at 6 weeks old were purchased from the Center for Laboratory Animals, Soochow University. After 1 week of acclimatization, diabetic mouse model was prepared as described previously with some modifications *(Lu et al., 2011; Li et al., 2017)*. The mice in normal group were fed with normal diet, and all the mice of other groups were fed with HFD (60% fat, 20% protein and 20% carbohydrate) during all the animal experiment. After 6 weeks of HFD feeding, the mice in the diabetic control model and imatinib group were fasted for 12 hours every night and injected with STZ (35 mg/kg, dissolved at 0.1 mM cold citrate buffer, pH 4.4) for 3 days to induce diabetes. Meanwhile, the control group was injected with citrate buffer. After a week of STZ injection, 12 h fasting glucose levels of all mice was measured. Mice with fasting blood glucose levels ≥ 11.1mmol/L *(Yu et al., 2014)* were considered as type 2 diabetes mice. Then, imatinib (40 mg/kg) was administered by intraperitoneal injection daily during the procedure. After 21 weeks injection of STZ, the mice were anesthetized with intraperitoneal injection of pentobarbital sodium. The hearts were dissected and stored at -80 ℃ for further analysis. During the experiment, mice were kept on their respective diets and their body weight was measured weekly. All procedures were performed with minimal damage to the mice.

### Immunofluorescence

After fixed in 4 % PFA and dehydrated with 20% sucrose, hearts were embedded in optimal cutting temperature (OCT) compound and stored at -80 °C. They were then sectioned by Leica CM1950 into 10µm-thick horizontal slices. The sections were incubated with primary antibody (anti-CD68 (Abcam, ab955), anti-CD31 (Abcam, ab28364), anti-CD31 (BD, 553700), anti-Pdgfb (CST, 3169T), anti-Pdgfc (Abcam, ab200401), anti-Pdgfd (Abcam, ab234666), anti-Pdgfra (R&D, AF1062-SP), anti-phospho-Pdgfra (Tyr754) (Thermo Fisher, 441008G), anti-Vim (R&D, BAM2105), anti-Hrc (Proteintech, 18142-1-AP), anti-Postn (R&D, AF2955-SP), anti-Itgb1 (Invitrogen, 14-0299-82), anti-FN1 (Abcam, ab2413), and anti-Lgals3bp (Abcam, ab236509)) or an IgG control for immunofluorescence staining. The fluorescent secondary antibodies (goat anti mouse IgM Alexa Fluor® 647, abcam, ab150123, or donkey anti rabbit IgG Alexa Fluor® 568, abcam, ab175470) and DAPI (SouthernBiotech, 0100-01) were used to visualize specific proteins.

### Real-time quantitative polymerase chain reaction (RT-qPCR)

Total RNA from mouse hearts was extracted using QIAGEN’s miRNeasy Mini kit (217004; Qiagen, Germany). The reverse transcription step was performed using Takara’s PrimeScriptTM RT Master Mix (RR036A; Takara, Japan). A brilliant SYBR green PCR master mix (4913914, Roche, Switzerland) was used to perform qPCR on cDNA templates in a LightCycler 480 (Roche, Switzerland). The target mRNA expression levels were normalized to that of GAPDH and the relative fold change was calculated using the 2^-ΔΔCT^ method. The qPCR primers for collagen I (forward 5’- AACTCCCTCCACCCCAATCT, reverse 5’-CCATGGAGATGCCAGATGGTT), collagen III (forward 5’–ACGTAAGCACTGGTGGACAG, reverse 5’– GGAGGGCCATAGCTGAACTG), Itgb1 (forward 5’-ATGCCAAATCTTGCGGAG AAT, reverse 5’-TTTGCTGCGATTGGTGACATT), and GAPDH (forward 5’– GGTCATCCATGACAACTT, reverse 5’–GGGGCCATCCACAGTCTT) are designed and synthesized by Invitrogen (Shanghai, China).

### siRNA-mediated knockdown of Itgb1 in mice heart

The siRNAs targeting Itgb1 were used as previously reported *(Speicher et al., 2014)*. Sense5’-AGAuGAGGuucAAuuuGAAdTsdT,antisense5’-UUcAAAUUGAACCUcA UCUdTsdT. Negative control: sense 5’-cuuAcGcuGAGuAcuucGAdTsdT, antisense 5’-UCGAAGuACUcAGCGuAAGdTsdT. These targeted siRNA sequences were subcloned into an AAV9 plasmid vector and packaged into AAV virus particle in vitro, and the titers of AAV viruses were ensured to exceed the 1x10^12^ vg/ml. Diabetic C57BL/6 male mice received negative control or Itgb1 siRNA mediated by AAV (0.5 mg/kg) via tail vein injection in a volume of 5 ml/kg body weight on the day 1 and 5 after the first STZ injection. At day 10 and month 5, the knockdown efficiency of Itgb1 was determined in the mice heart.

### Isolation of nuclei from heart tissue

Isolation of nuclei from heart tissue were analyzed as previously described *(McLellan et al., 2020)*. Briefly, mouse heart tissues were homogenized using a Wheaton Dounce Tissue Grinder. 3 ml of homogenization buffer was added and the homogenized tissue was incubated on ice for 5 minutes. Then the homogenized tissue was filtered through a through 40 mm cell strainer, mixed with an equal volume of working solution and loaded on top of an OptiPrep density gradient on top of 5 ml 35% OptiPrep solution. The nuclei were separated by ultracentrifugation using an SW32 rotor (20 minutes, 9000 rpm). 3 ml of nuclei were collected from the 29%/35% interphase and washed with 30 ml of PBS containing 0.04% BSA. The nuclei were centrifuged at 300 g for 3 minutes and washed with 20 ml of PBS containing 0.04% BSA. Then the nuclei were centrifuged at 300 g for 3 minutes and re-suspended in 500 microliter PBS containing 0.04% BSA. All procedures were carried out on ice or at 4 °C.

### Single-nucleus transcriptomic library preparation

Single-nucleus transcriptomic library preparation were performed as previously described *(Li et al., 2021)*. Briefly, single nucleus was resuspended in PBS with 0.04% BSA and added to each channel. The captured nucleus was lysed, and the released RNA was barcoded through reverse transcription in individual GEMs. Barcoded cDNA was amplified, and the quality was controlled using Agilent 4200 TapeStation System. scRNA-seq libraries were prepared using Single Cell 3′ Library and Gel Bead Kit V3 following the manufacture’s introduction. Sequencing was performed on an Illumina Novaseq 6000 sequencer with a pair-end 150 bp (PE150) reading strategy (performed by Gene Denovo Biotechnology Co., Guangzhou, China).

### Clustering analysis

Alignment, filtering, barcode counting, and UMI counting were performed with Cell Ranger to generate a feature-barcode matrix and their global gene expressions. Dimensionality reduction, visualization, and analysis of scRNA-sequencing data were performed with the R package Seurat (version 3.1.2). As a further quality-control measure, nucleus meeting any of the following criteria were filtered out: <500 or >4,000 unique genes expressed, >8,000 UMIs, or >10% of reads mapping to mitochondria. After removing unwanted nucleus from the dataset, two thousand highly variable genes were used for downstream clustering analysis. Principal Component Analysis (PCA) was performed, and the number of the significant principal components was calculated using the built-in “ElbowPlot” function.

### Differentially expressed genes analysis

Expression level of each gene in target cluster were compared against the rest of cells using Wilcoxon rank sum test. Significant upregulated genes were identified using the following criteria: (i) at least 1.28-fold overexpressed in the target cluster, (ii) expressed in more than 25% of the cells belonging to the target cluster, and (iii) FDR is less than 0.05.

### Gene Set Variation Analysis

To identify cellular processes and pathways enriched in different clusters, Gene Set Variation Analysis (GSVA) was performed in the GSVA R package *(Hänzelmann et al., 2013)* version 1.26 based on the cluster-averaged log-transformed expression matrix.

### Differentially expressed genes Gene Ontology and Kyoto Encyclopedia of Genes and Genomes pathway enrichment analysis

Gene Ontology (GO) and Kyoto Encyclopedia of Genes and Genomes (KEGG) pathway enrichment analysis identified significantly enriched cellular processes and pathways in differentially expressed genes comparing with the whole genome background. The calculated p-value was gone through FDR Correction, taking FDR ≤ 0.05 as a threshold. GO term and KEGG pathways meeting this condition were defined as significantly enriched pathways in differentially expressed genes.

### Regulon analysis

Regulon analysis was performed on the SCENIC R package to carry out transcription factor network inference *(Aibar et al., 2017)*. In brief, gene expression matrix was used as input, and the pipeline was implanted in three steps. First, gene co-expression network was identified via GENIE3 *(Huynh-Thu et al., 2010)*. Second, we pruned each module based on a regulatory motif near a transcription start site via RcisTarget. Third, we scored the activity of each regulon for each single cell via the AUC scores using AUCell R package.

### Statistical analyses

The statistical analysis of the results was performed using GraphPad Prism 9.0 software. Unpaired t test or one-way ANOVA analysis were used to calculate the differences in mean values. P ≤ 0.05 was considered statistically significant. Other statistical analyses not described above were performed using the ggpubr package in R (https://github.com/kassambara/ggpubr).

## Acknowledgments

We thank Guangzhou Gene Denovo Biotechnology Co. (Guangzhou, China) for the technical support in single cell RNA sequencing of normal and T2DM mouse heart, and thank Professor Huiling Zhang’s team of College of Pharmaceutical Science in Soochow University for their friendly help in the experimental process of diabetic myocardial fibrosis in mice. Thanks to the public research platform of Soochow University for their technical support of this experiment. We also thanks to the native English speaker from CureEdit to improve the grammar and readability.

---

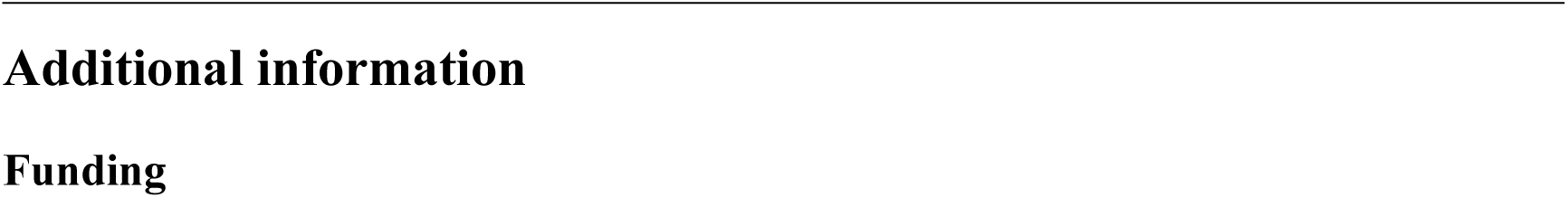
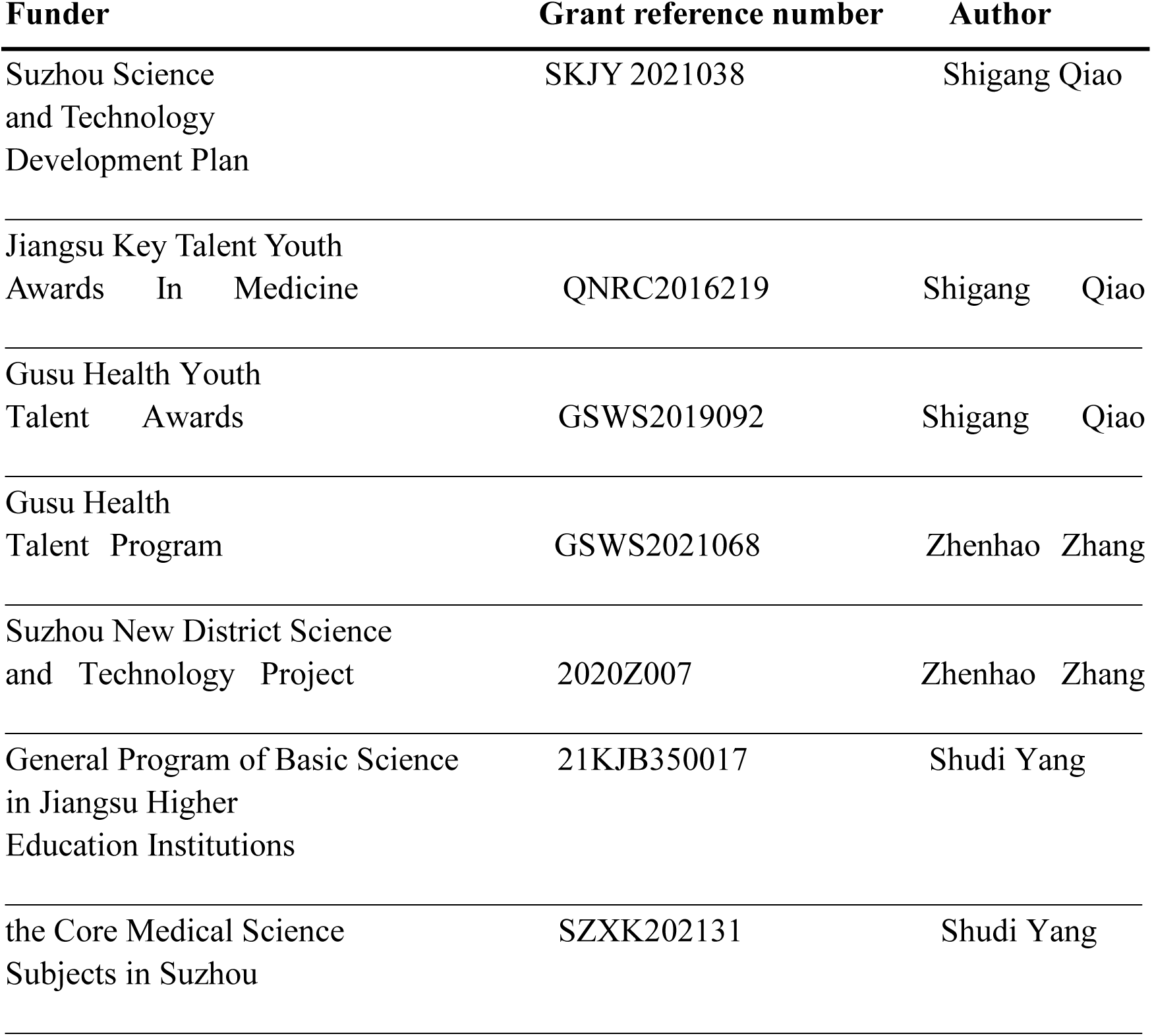

## Author Contributions

Wei Li, Conceptualization, Data curation, Formal analysis, Methodology, Investigation, Writing - original draft, Writing - review and editing; Xinqi Lou, Yingjie Zha, Jun Zha, Lei Hong, Shudi Yang and Zhanli Xie, Formal analysis, Investigation; Chen Wang, Conceptualization; Jianzhong An, Conceptualization, Guidance; Zhenhao Zhang, Conceptualization, Funding acquisition, Writing - original draft, Writing - review and editing; Shigang Qiao, Conceptualization, Funding acquisition, Supervision, Writing - original draft, Writing - review and editing.

## Author ORCIDs

Zhenhao Zhang https://orcid.org/0000-0001-5659-6315

Shigang Qiao https://orcid.org/0000-0003-0309-8405

## Ethics

Our present study was approved by the ethics committee of Soochow University and Suzhou Science & Technology Town Hospital, Gusu School, Nanjing Medical University. All mouse were treated in accordance with the National Institutes of Health’s Guidelines for the Care and Use of Experimental Animals (NIH publications No. 80-23, revised 1996).

## Additional Files

### Supplementary files

Supplementary file 1: 25 transcriptionally distinct pre-clusters with highly consistent expression patterns across individual mouse heart.

Supplementary file 2: Genes of significant transcriptomic changes in cardiac populations.

Supplementary file 3: The top 10 upregulated genes in cardiac populations. Supplementary file 4: unique differentially-expressed genes (uni-DEGs) in cardiac populations.

Supplementary file 5: Significantly differentially-expressed genes in specific cell populations relative to others in mouse heart.

Supplementary file 6: Cell type-specific receptors in cardiac populations.

Supplementary file 7: Cell type-specific ligands in cardiac populations.

Supplementary file 8: Relative expression of a selection of essential growth factors across major cardiac cell types.

Supplementary file 9: The number of ligand-receptor pairs between cardiac cell populations in healthy mice.

Supplementary file 10: Ligand-receptor pairs between cardiac cell populations in healthy mice.

Supplementary file 11: Significant differentially-expressed ligands for each cell population.

Supplementary file 12: Significant differentially-expressed receptors for each cell population.

Supplementary file 13: The cognate ligands of Egfr. Supplementary file 14: The cognate ligands of Pdgfra.

Supplementary file 15: 10 transcriptionally distinct fibroblast subpopulations.

Supplementary file 16: Distinct signatures of each fibroblast subpopulations in heart.

Supplementary file 17: The transcription-factor gene-regulatory networks in the distinct subpopulations.

Supplementary file 18: Gene-expression of transcription factors in Figure 5A and B.

Supplementary file 19: Genes of significant transcriptomic changes in each fibroblast subpopulation.

Supplementary file 20: unique differentially-expressed genes (uni-DEGs) in each fibroblast subpopulation.

Supplementary file 21: The cognate ligands of Itgb1. Transparent reporting form

## Data availability

All sequencing data that support this study is available at Genome Sequence Archive in BIG Data Center (http://bigd.big.ac.cn/) with the accession code CRA007245. Ligand and receptor pairing dataset was obtained from Fantom5 (https://fantom.gsc.riken.jp/5/suppl/Ramilowski_et_al_2015/), as recently described (Ramilowski et al., 2015). SNPs of Itgb1 and Fn1 correlated with type 2 diabetes and glucose metabolic disorders generated in this study was obtained from GWAS CENTRAL (https://www.gwascentral.org/). Source data files are provided to support CT values of Collagen I and Collagen III used for Figure 3G-H. Source data files are provided to support CT values of Itgb1 used for Figure 6H.

The following dataset was generated:

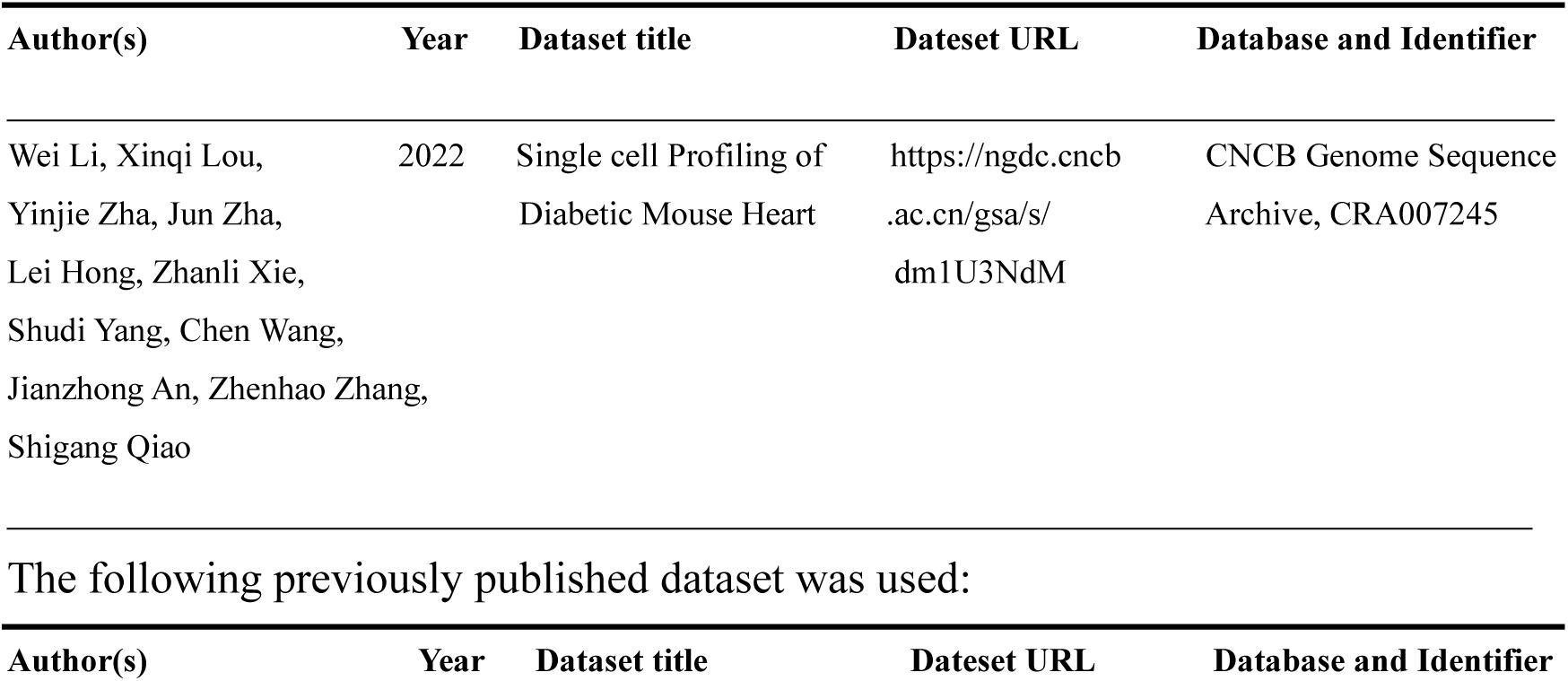
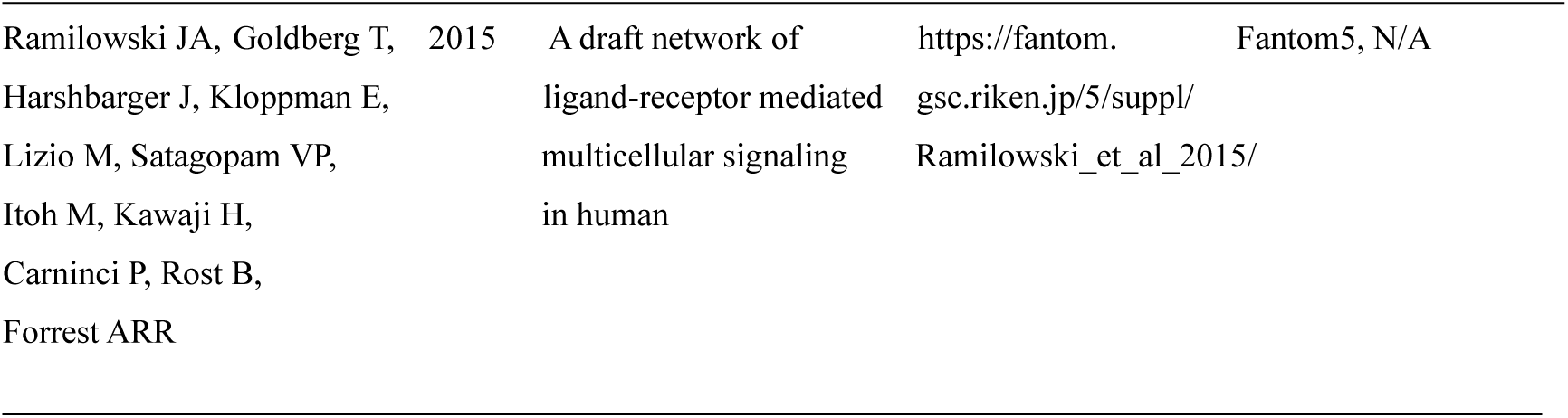

